# Biosynthetic plasticity enables CD8+ T cell functional resilience under nutrient stress

**DOI:** 10.1101/2025.01.22.632829

**Authors:** Michael Scaglione, Montana Knight, Krittin Trihemasava, Kelly Rome, Anne-Sophie Archambault, Juhee Oh, Erin Tanaka, Elise Hall, Tran Ngoc Van Le, Caleb L. Lines, Brian Goldspiel, Hossein Fazelinia, Clemence Queriault, Lucien Turner, Tanay Parnaik, Jimmy Xu, Lynn A. Spruce, Caroline Bartman, Clementina Mesaros, Ramon I. Klein Geltink, Crystal S. Conn, Will Bailis

## Abstract

To maintain lineage-specific functions, cells must acquire and allocate nutrients across diverse cellular processes, even in metabolically-dysregulated environments. The mechanisms allowing CD8+ T cells to maintain immune function in perturbed environments are poorly understood. We find that CD8+ T cells adapt to nutrient stresses over time, reconfiguring gene-regulatory and metabolic networks to license functional recovery. Under acute stress, T cells reorient translational programming, limiting nutrient demand while prioritizing stress-sensitive metabolic and transcriptional responses. Within these responses, the transcription factors ATF4 and CEBPG jointly establish an adaptive metabolic program, promoting amino acid synthesis and uptake while maintaining mitochondrial anaplerosis. Despite diminished energetic capacity under environmental stress, this program prevents failure of central carbon metabolism, mitigating stress amplification and cellular dysfunction to potentiate anti-tumor immunity. Altogether, we demonstrate that biosynthetic plasticity via translational and metabolic reprioritization confers functional resilience to immune cells in unfavorable environments, offering novel strategies to enhance immunotherapies.

## Introduction

Individual cells within multicellular organisms require nutrients to support fundamental cellular processes (e.g. growth, viability) as well as lineage-specific functions that promote organismal health. To do this, cells must acquire adequate nutrients from the environment and coordinate their utilization by a variety of competing biochemical pathways, such as protein translation, energy generation, or antioxidant synthesis^1,2^. As both cell homeostasis and lineage-specific functionality are dependent on a common store of intracellular nutrients, cells must cohesively regulate global nutrient consumption and pathway-level allocation, integrating information from growth signaling, functional programing, and environmental nutrient availability. While there is an appreciation for how malignant cells favor nutrient use for cell survival and proliferation at the expense of the host, how healthy non-transformed cells harmonize the metabolic requirements of host-protective functionality and fundamental cellular homeostasis remains poorly understood.

To ensure that metabolic demand and nutrient supply are continuously balanced, cells employ diverse strategies to sense intracellular and environmental metabolic state and couple this information to the regulatory control of global biosynthesis and nutrient handling^3–5^. For example, protein translation, a primary consumer of energy and amino acids, is a key hub for regulatory control through nutrient-sensitive signaling pathways including the mechanistic target of rapamycin (mTOR) and the integrated stress response (ISR) pathways^6–10^. These signaling pathways also influence metabolic activity, nutrient acquisition, and regulatory responses, broadly coupling cell biology to environmental state. Although these conserved pathways control suites of cellular pathways across cell lineages, how these processes intersect with lineage-specific programming to control cell function is an active area of investigation. Additionally, how cells dynamically remodel highly interconnected metabolic, regulatory, and functional networks in response to changing or limiting environmental conditions is currently underexplored^11–15^.

The ability to adapt to dynamic environments is integral to the biology of circulatory immune cells, including T cells. During an immune response, circulating CD8+ T cells travel to perturbed peripheral sites (e.g. tumors or infected tissue) to perform protective effector functions such as cytokine production or cytotoxicity. These sites are biochemically distinct from blood, containing varying levels of nutrients as well as inhibitory or immunosuppressive metabolites^15–18^. Indeed, nutrient deprivation and metabolic dysregulation are appreciated to drive cellular dysfunction and limit cellular persistence^19–29^. Despite this, there is limited insight into factors that mitigate stress-driven dysfunction and support functional resilience — continued immune function in changing or limiting environmental conditions. Additionally, though there is a growing appreciation that metabolic perturbation and sensing can regulate gene expression and immune cell function, the global regulatory impact, temporal dynamics, and specificity of nutrient-stress-induced changes in T cell programming are poorly understood^30–39^. Understanding how immune cells adapt to and overcome environmental stress would offer novel axes of immunotherapeutic intervention^40^.

Here, we set out to illuminate the global regulatory underpinnings of nutrient stress adaptation in CD8+ T cells, aiming to identify factors that promote immune function in limiting or stressful metabolic environments. We show that despite an initial loss of immune function upon acute nutrient stress, CD8+ T cells exhibit the capacity for metabolic adaptation and functional recovery over time. Using multi-omic analyses of a diverse set of nutrient stresses, we comprehensively resolve stress-sensitive programming in CD8+ T cells, identifying nutrient-sensitive molecular targets within the transcriptome, translatome, and proteome and revealing the translatome to be particularly dynamic. We find that early during periods of nutrient stress, initial loss of mTORC1 signaling promotes the translational repression of mitochondrial and ribosomal mRNAs, preserving amino acid pools by reducing metabolic demand. Concurrently, activation of ISR signaling induces a stress-sensitive transcriptional network to drive amino acid synthesis and transport, promoting metabolic adaptation and recovery of immune function. We highlight two stress-induced transcription factors, ATF4 and CEBPG, as key mediators of adaptive metabolic gene expression. Despite diminished energetic capacity under environmental stress, these factors act in concert to promote amino acid synthesis and uptake and maintain mitochondrial anaplerosis to promote continued immune function, revealing a novel metabolic mechanism that mitigates stress-driven cellular dysfunction to potentiate anti-tumor immunity. Together, these data illustrate how CD8+ T cells concordantly reprogram global gene regulation and metabolism to maintain immune function in the face of stress, providing insight into environmental control of immune function for immunotherapeutic applications as well as a paradigm for broader inquiry into resilience of lineage-specific cell functions across biochemical environments.

## Results

### Nutrient stress drives metabolic and gene-regulatory adaptation in CD8+ T cells to support recovery of immune function

To assess the ability for effector-like CD8+ T cells to adapt to nutrient stress over time, we assayed the metabolic content and functional capacity of previously-activated CD8+ T cells during an acute (6 hour) or long-term (24 hour) exposure to culture medium conditioned by the MC38 colorectal cancer cell line (“tumor supernatant”) (Fig 1A). This conditioned medium was depleted of many amino acids – including glutamate, methionine, and serine – and enriched for proline and alanine relative to non-conditioned medium (Fig 1B). After 6 hours of culture in tumor supernatant, CD8+ T cells showed an altered intracellular amino acid profile that largely mirrored the content of the medium, and cells restimulated during this time showed decreased cytokine production capacity relative to cells restimulated in control medium (Fig 1 C and D, Fig S1A). However, by 24 hours of exposure to tumor supernatant, the intracellular levels of many previously-depleted amino acids recovered, and CD8+ T cells restimulated at this time showed rescued or even increased functional capacity relative to control medium with no loss in viability (Fig 1 C and D, Fig S1A). Moreover, we observed an increase in the surface expression of CD98, a common heavy chain subunit of System L and System Xc-amino acid transporters when dimerized with various light chains including SLC7A5 (LAT-1) and SLC7A11 (xCT) (Fig 1E)^41^. We also observed an increase in the mRNAs for Slc3a2, Slc7a5, and Slc7a11 within 6 hours of culture in tumor supernatant (Fig S1B). Together, these data suggest that despite sensitivity to acute changes in metabolic environment, CD8+ T cells retain an ability to rebalance their intracellular metabolic state and restore functional capacity over time, correlating with increased expression of amino acid transporters such as LAT-1 and xCT.

**Figure 1:**
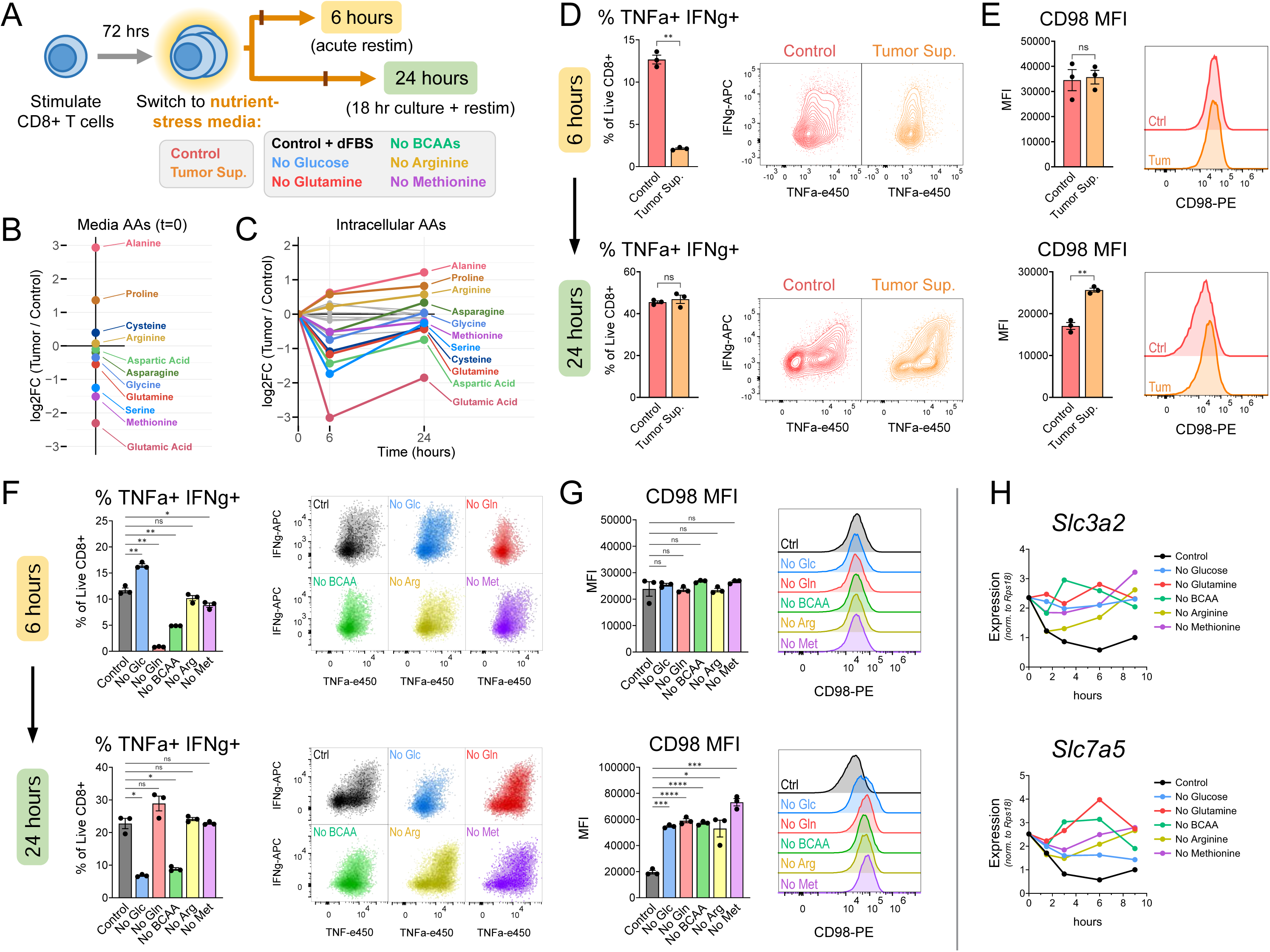
CD8+ T cells exhibit genetic and metabolic reprogramming during nutrient stress adaptation. A) Schematic of nutrient stress models. Previously-activated mouse CD8+ T cells were treated with tumor supernatant or nutrient-depleted media for a total of 6 or 24 hours and re-stimulated during the final 6 hours of culture. B) Relative difference in medium amino acid abundance in tumor-conditioned medium vs. control medium. C) Relative difference in cellular amino acid abundance in CD8+ T cells treated with tumor-conditioned medium vs. control medium after 6 or 24 hours of culture. Amino acids increased over time are shown in color. D-E) Intracellular cytokine production (D) and surface CD98 expression (E) in CD8+ T cells treated with tumor-conditioned medium for 6 hours (top) or 24 hours (bottom) F-G) Intracellular cytokine production (F) and surface CD98 expression (G) in CD8+ T cells treated with nutrient stress media for 6 hours (top) or 24 hours (bottom) H) Expression of amino acid transporter transcripts over time in CD8+ T cells treated with nutrient stress media

To better understand the extent and specificity of their stress-adaptation capabilities, we assayed the function of CD8+ T cells over time using an array of nutrient-depleted media formulated without either glucose, glutamine, branched-chain amino acids (BCAAs – leucine, isoleucine, valine), arginine, or methionine (Fig 1A). These metabolites were chosen as they have broad differences in essentiality, utilization by cellular catabolism and biosynthesis pathways, and sensing by signaling pathways^42^. We confirmed that each respective amino acid removed from our nutrient-stress media was substantially depleted from intracellular pools within 3-6 hours, again suggesting rapid equilibration of intracellular amino acid pools with the extracellular environment (Fig S1C). CD8+ T cells cultured in these nutrient-stress media exhibited marked diversity in their acute and long-term outcomes depending on the medium composition: cells cultured without glutamine were able to completely rescue early deficits in function over time (similar to those treated with tumor supernatant), cells cultured without BCAAs retained full viability but were unable to recover immune function, cells cultured without glucose lost immune function and eventually underwent cell death, and cells cultured without arginine or methionine displayed minimal changes in immune function over time (Fig 1F, Fig S1D). In spite of this, cells in each condition displayed a marked increase in surface CD98 expression over 24 hours, with increases in Slc3a2 and Slc7a5 transcripts detectable within hours (Fig 1G, Fig 1H). These data reveal that T cells differ in the quality and magnitude of their response to individual nutrients as well as their ability to overcome the complete absence of these nutrients over time. Despite this, CD8+ T cells rapidly initiate a conserved amino acid uptake response to a diverse array of nutrient stresses and this response is correlated with metabolic and functional adaptation within particular nutrient-depleted environments (Fig S1E).

### Acute nutrient stress reorients nutrient allocation by limiting the translation of a biosynthetic program to favor stress adaptation programming

Next, we aimed to better understand the acute response to nutrient stress at a global level, hypothesizing that when nutrients are limited, cells may modify current gene-regulatory networks to minimize energy expenditure and nutrient usage while engaging processes to restore cellular homeostasis and immune function. As cells can modify gene expression through transcriptional, translational, and post-translational mechanisms, we tested which pool would be most sensitive to stress, which particular targets within each pool would be most dynamic, and whether these responses would be unique or conserved across various nutrient stress models^43^.

To comprehensively dissect the breadth and specificity of acute nutrient stress responses in CD8+ T cells, we profiled nutrient-stress-sensitive programming using total RNA sequencing, polysome-fractionation, and whole-cell proteomics after 3 hours of culture in five different nutrient stress media (Fig 2A). Each condition dampened global translation as shown by a decrease in polysome abundance and polysome/monosome ratio (Fig 2B and C). Analysis of the transcriptome and translatome composition revealed a variety of protein coding and non-coding transcripts, including several long-non-coding RNAs (Fig S2A-C). While patterns of relative transcript abundance between the transcriptome and translatome were distinct, we observed that acute nutrient stress drove rapid remodeling of both pools, with the polysome-bound RNA responses revealing the greatest number of differentially-expressed transcripts and the greatest magnitude of stress-driven changes (Fig 2D, Fig S2D-F). On the other hand, changes in the proteome were limited after 3 hours of nutrient stress, highlighting the rapid nature of RNA responses to stress (Fig 2D).

**Figure 2:**
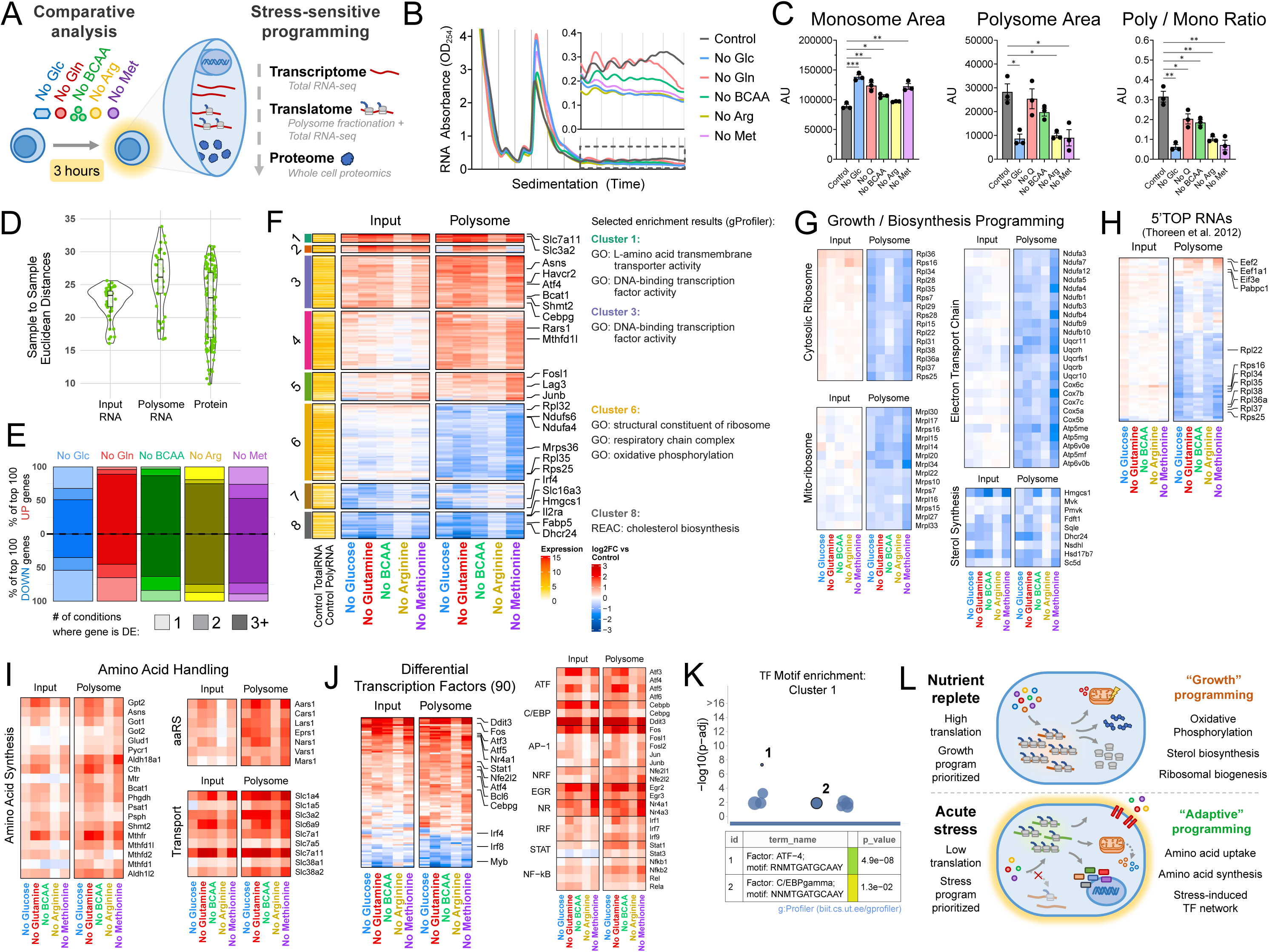
Acute nutrient stress drives translational reprioritization of pro-adaptive programming over growth programming. A) Schematic of comparative analysis of the stress-sensitive programming (transcriptome, translatome, and proteome) in CD8+ T cells. Cells were treated with nutrient stress media for 3 hours before performing polysome fractionation, total RNA sequencing of input and polysome-bound RNA, and whole-cell proteomics B) Polysome profiles of CD8+ T cells after 3 hours of culture in nutrient stress conditions. Inset shows polysome region. C) Quantification of monosome area, polysome area, and polysome / monosome ratio from polysome profiles. D) Plot of control-treatment sample-sample Euclidean distances in input RNA, polysome RNA, and protein datasets. E) Number of differentially expressed genes within the top 100 DEGs (up or down) found to be uniquely differential to each condition (lightest), or differential in the same direction across 2 (middle) or 3+ (dark) total nutrient stress conditions F) Global heatmap of differentially expressed transcripts in input RNA (left) and polysome-associated RNA (right) in nutrient stress conditions relative to control. Left annotation: k-means cluster membership and relative expression in control condition. Right annotation: selected pathways enriched in indicated clusters G) Change in expression of growth/biosynthesis-related transcripts in input RNA (left) and polysome-associated RNA (right) in nutrient stress conditions H) Change in expression of all transcripts containing a 5’ TOP motif from Thoreen et al. (2012) in input (left) and polysome-associated RNA (right) I) Change in expression of transcripts related to amino acid handling in input RNA (left) and polysome-associated RNA (right) in nutrient stress conditions. J) Change in expression of all differential transcription factors and selected stress-related transcription factors in input RNA (left) and polysome-associated RNA (right) in nutrient stress conditions. K) Motif enrichment results for transcripts in Cluster 1 L) Schematic for translational reprioritization during acute nutrient stress

We observed that stress-driven changes in both the transcriptome and translatome were largely conserved across nutrient depletion conditions; many differential transcripts trended in the same direction in multiple stress conditions, and a majority of the top 100 differentially-expressed transcripts in the polysome-bound pool were differentially expressed in the same direction in at least 3 total conditions (Fig 2E and F, Fig S2G). However, we also found nutrient-specific modules in each response, with glucose depletion and methionine depletion generating the most unique signatures (Fig 2E and F, Fig S2G). Lastly, we saw that the magnitude of each stress-response differed between conditions. Whereas arginine depletion produced the fewest differentially-expressed transcripts and weakest changes in transcript abundance, methionine depletion produced the most differentially-expressed transcripts and the greatest magnitude of response (Fig 2F, Fig S2G). Taken together, these data suggest that nutrient depletion drives rapid remodeling of the transcriptome and translatome, revealing translation to be particularly responsive to nutrient stress. Although each condition displays specificity with regard to the subsets of transcripts affected and the magnitude of the overall effect, these acute RNA responses are largely conserved across diverse conditions, highlighting a core stress-response program.

Next, we aimed to identify the major stress-sensitive pathways and regulatory networks altered upon acute nutrient stress. We noted that key transcripts associated with effector T cell function (Gzmb, Ifng, Il2ra, and glycolysis) were among the most abundant in the translatome in control conditions and that the relative prioritization of these transcripts was maintained even after nutrient stress, in contrast to other abundant transcripts such as those encoding translational machinery s, suggestive of selective regulation of cell programming (Fig S3F). We then focused our analysis on highly differential pathways as well as pathways that were differentially affected between the transcriptome and translatome, suggestive of translational prioritization or repression under stress. To do so, we performed k-means clustering on the stress-driven changes in gene expression in both the input and polysome-associated RNA samples and subjected these clusters to pathway analysis (Fig 2F). While some clusters showed similar trends in stress-driven changes between input and polysome-bound RNA samples, other clusters displayed stronger responses within the polysome-bound RNA samples, suggestive of translational prioritization or repression under stress (Fig 2F, Fig S3A and B). Of note, Cluster 6 contained transcripts that were disproportionately depleted from the polysome-bound pool, indicating translational repression. This cluster was enriched for transcripts related to ribosomal and mitochondrial biology, including cytosolic ribosomes, mitoribosomes, and oxidative phosphorylation (OXPHOS) (Fig 2F and G). These changes were robust at the pathway level, with a majority of genes in each pathway exhibiting a similar depletion in the polysome-associated RNA pool relative to input RNA in all five nutrient stresses (Fig S2H). We noted that many of the transcripts in this cluster contain a 5’ TOP motif, an RNA motif allowing for translational regulation downstream of mTOR signaling that has been previously implicated in effector T cell expansion and contraction (Fig 2H)^44–49^. Moreover, other transcripts related to cell growth, including those involved in cholesterol biosynthesis (Hmgcs1, Mvk, Dhcr24, Hsd17b7 and Scd5) and growth factor signaling (Il2ra), were also depleted from both the input RNA and polysome-bound RNA pools across all conditions (Fig 2F and G). Together, these data suggest nutrient stress drives CD8+ T cells to deprioritize a gene regulatory program related to growth and global biosynthesis.

In addition to transcripts with decreased expression, we observed multiple clusters containing transcripts with elevated expression that were often disproportionately enriched in the polysome-bound RNA pool under nutrient stress, suggestive transcriptional activation and/or translational prioritization (Fig 2F). These clusters were enriched for transcripts related to amino acid handling, including those related to tRNA charging (*Lars1, Cars1, Aars1),* amino acid transport (*Slc7a5, Slc3a2, Slc6a9, Slc1a4, Slc7a11* etc.) and amino acid synthesis, particularly for alanine (*Gpt2),* proline (*Pycr1* and *Aldh18a1)*, serine/glycine (*Phgdh, Psat1, Psph, Shmt2 etc.)*, and asparagine (*Asns*) (Fig 2F and I). These responses were observed across all conditions, suggesting that CD8+ T cells rapidly induce a conserved metabolic program during the acute response to nutrient stress, without regard for nutrient identity. Stressed T cells also displayed increased *Havcr2* (encoding Tim3) and *Lag3* transcript levels, suggesting a potential link between nutrient stress and immunoregulation via expression of inhibitory receptors (Fig 2F). Additionally, many transcripts containing internal ribosomal entry sites (IRES) were enriched in polysome-associated RNA samples relative to input RNA, suggesting that the transcripts that are prioritized during stress may contain unique regulatory features that underlie their continued association with polysomes (Fig S2I).

Lastly, we noted that an array of 90 transcription factors were acutely sensitive to nutrient stress, as defined by being differentially expressed in at least one stress condition relative to control. A majority of these factors were upregulated, including members of the ATF (*Atf3, Atf4, Atf5, Atf6*), CEBP (*Cebpb, Cebpg*), NRF (*Nfe2l1, Nfe2l2*), and AP-1 (*Fos, Jun)* families, among others (Fig 2J). We also observed that putative transcription factor binding sites for a subset of these factors (including ATF4 and CEBPG) were enriched near the transcriptional start sites of transcripts that were most strongly induced under stress (Cluster 1), suggesting that these factors play an outsized role in mediating the early response to nutrient stress in CD8+ T cells (Fig 2K).

While the proteome of nutrient-stressed T cells was less dynamic, we observed several differentially expressed proteins upon nutrient stress. These tended to correlate between changes in protein abundance and polysome-bound RNA levels, including *Egr2, Ifrd1, Txnip, Irf8, Slc6a9, Fos,* and *Il2ra* (Fig S3C). Taken together, this suggests that although the RNA dynamics occur rapidly, some important stress-sensitive RNA responses are detectable within the proteome during this acute response.

Taken together, our data suggest that CD8+ T cells undergo a rapid remodeling of gene expression networks across the transcriptome, translatome, and proteome in response to nutrient stress, with marked conservation in RNA responses across conditions. Under acute stress, cells decrease the translational priority of growth-related gene expression while elevating expression of stress-adaptive programming, including a suite of genes involved in amino acid uptake and synthesis (Fig 2L). These regulatory changes may act to divert nutrient allocation away from biomass generation, instead promoting the translation of programming that assists the cell in restoring metabolic homeostasis and immune function. Lastly, stress-responsive transcription factors such as ATF4 and CEBPG appear to have an outsized role in mediating these early transcriptional responses, highlighting a potential link between regulatory and metabolic reprogramming and recovery of immune function in nutrient stressed CD8+ T cells.

### Loss of mTORC1 / translation limits metabolic demand, while ISR signaling drives amino acid programming to restore immune function

We next aimed to dissect the upstream pathways leading to these transcriptional and translational responses. Both mTOR and ISR signaling are known to broadly regulate metabolism, gene expression, and cap-dependent translation in response to nutrient availability or cellular stressors^47,50^. Consistent with this, we observed that nutrient stress broadly dampened mTORC1 activity and activated the ISR, as evidenced by decreased phosphorylation of small ribosomal protein subunit 6 (p-S6) and increased ATF4 levels, respectively (Fig 3A, Fig S4A). We further observed that nutrient stress rapidly induced a broad set of ISR-related transcripts, including Atf3, Nupr1, Ppp1r15a, and Sesn2 (Fig 3B, Fig S4B)^51^. Notably, this gene signature was also induced across a range of nutrient concentrations, suggesting that the magnitude of ISR signaling in CD8+ T cells scales with nutrient abundance (Fig S4C). Similarly, we observed dampened p-S6, increased ATF4 levels, and induction of stress-related transcripts when cells were cultured in tumor supernatant, highlighting the conserved nature of these responses (Fig S4D, Fig S4E).

**Figure 3:**
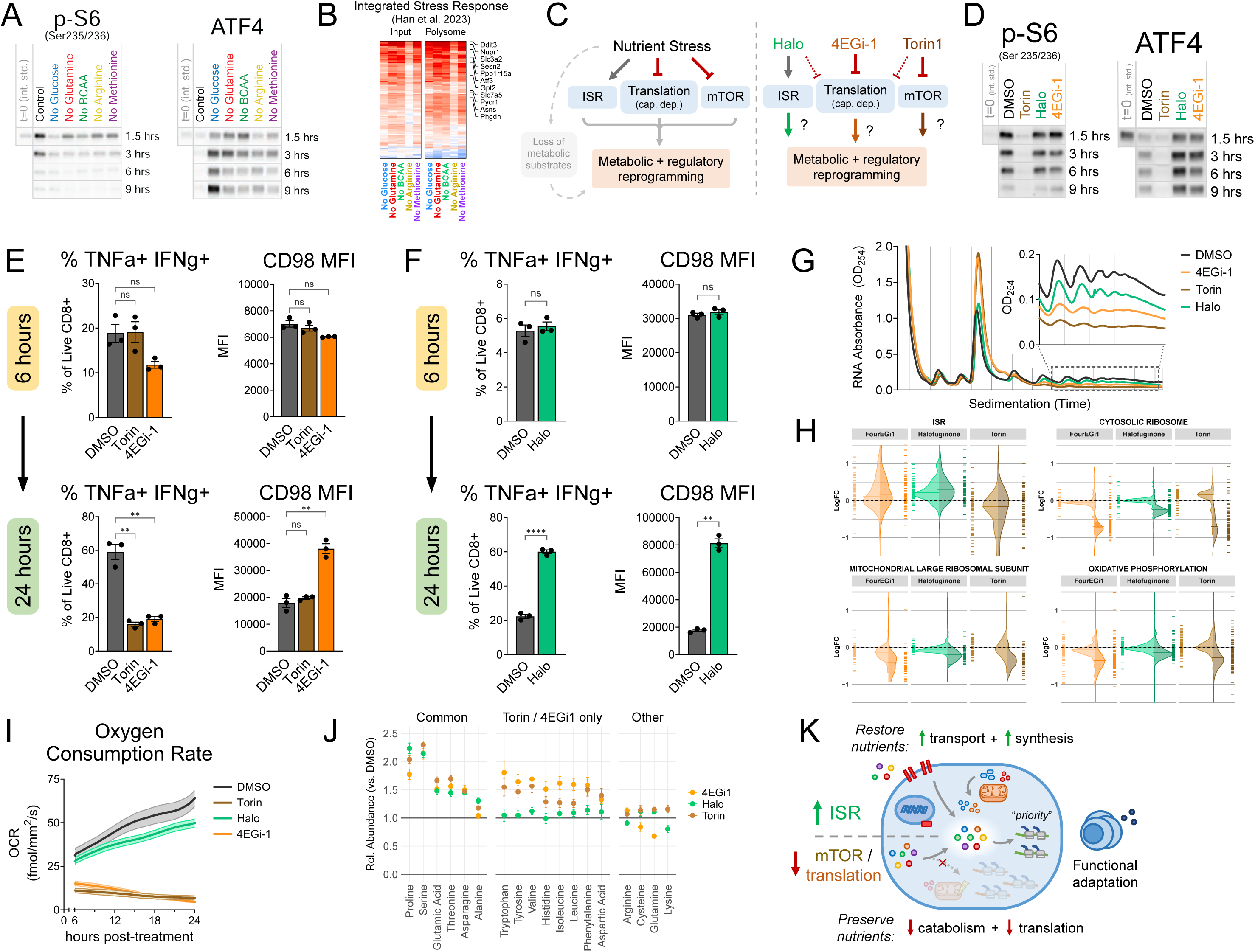
The ISR, mTOR, and global translational capacity govern discrete metabolic and regulatory adaptations. A) Visualization of mTORC1 signaling (p-S6) and ISR signaling (ATF4) over time in CD8+ T cells cultured in nutrient stress media B) Heatmap of change in ISR-related transcripts from Han et al. 2023 in nutrient stress media vs control. C) Schematic of approach to deconvolute effects of stress-induced signaling and translation on gene-regulatory and metabolic reprogramming observed in CD8+ T cells under nutrient stress. Left: Nutrient stress models concurrently alter nutrient-sensitive signaling and global translation while decreasing metabolite availability in the environment. Right: Pharmacological tools individually replicate stress-driven changes in signaling and translation to isolate the contribution of each pathway in reprogramming gene regulation and metabolism without confounding metabolic effects from changes in environmental substrate abundance. Torin1 = mTOR inhibitor; Halo = halofuginone, inhibitor of Eprs, activates ISR via GCN2; 4EGi-1 = inhibitor of cap-dependent translation. D) Effect of drug treatments on mTORC1 signaling (p-S6) and ISR signaling (ATF4) over time. E) Intracellular cytokine production and CD98 expression in CD8+ T cells treated with Torin or 4EGi1 for 6 hours (top) or 24 hours (bottom). F) Intracellular cytokine production and CD98 expression in CD8+ T cells treated with halofuginone for 6 hours (top) or 24 hours (bottom). G) Polysome profiles of CD8+ T cells treated with drug treatments for 3 hours. Inset shows data for polysome region. H) Change in ISR related transcripts and ribosomal/mitochondrial pathways in drug treatments (3 hours) vs control for input RNA (left), and polysome-associated RNA (right) I) Oxygen consumption rate of cells treated with drug treatments over time via Resipher assay J) Relative cellular amino acid abundance under drug treatments at 6 hours (vs. DMSO). K) Model of the contributions of ISR / mTOR / global translation to nutrient handling and metabolic programming during stress adaptation

Having observed the concurrent loss of mTORC1 signaling, induction of the ISR, and suppression of global translation during acute nutrient stress, we aimed to dissect the individual contribution of these stress-driven phenotypes to the gene-regulatory, metabolic, and functional responses we observed as CD8+ T cells adapted to nutrient stress. As the nutrient depletion models affect stress-sensitive gene regulation as well as impose limitations on metabolic substrate availability for translation or energy production, we opted to use pharmacological methods to model the regulatory, metabolic, and functional effects of these stress-sensitive signaling networks and translation in isolation without perturbing nutrient availability (Fig 3C).

To this end, we cultured activated CD8+ T cells with either Torin1 (an mTOR inhibitor), halofuginone (an inhibitor of glutamyl-prolyl tRNA synthetase, leading to ISR activation), or 4EGi-1 (an inhibitor of eIF4E-eIF4G interaction, inhibiting eIF4F-dependent translation) (Fig 3C)^52–54^. As expected, we observed a loss of p-S6 upon Torin1 treatment, while halofuginone-treated cells displayed increased ATF4 levels and an induction of stress-related transcripts (Fig 3D, Fig S5A and B). We further noted that ATF4 levels and several stress-related transcripts were decreased below control levels when cells were treated with Torin, in line with reports in other cell types that mTORC1 activity can drive ATF4-dependent gene expression and metabolic activity in the absence of extrinsic stress^55–59^. Treatment with 4EGi-1 was also sufficient to drive an increase in ATF4 protein and stress-related transcript expression, suggesting translational insults alone may be sufficient to drive ISR induction (Fig 3D, Fig S5A and B).

We then evaluated how each drug affected T cell function over time. We found that loss of mTOR signaling or dampening global cap-dependent translation with 4EGi-1 treatment led to an acute loss of cytokine production that failed to recover over 24 hours (Fig 3E, Fig S5C). Conversely, ISR activation did not interfere with short-term immune function and notably, CD8+ T cells displayed a marked increase in effector function following prolonged ISR signaling (Fig 3F, Fig S5D and E). ISR activation led to rapid induction of Slc7a5 and Slc3a2 transcripts and CD98 protein expression over 24 hours, highlighting ISR signaling as a key driver of stress-induced CD98 expression in CD8+ T cells, in accordance with previous studies in cancer cells (Fig 3F, Fig S5E, and F)^60^. Alongside the functional recovery observed in the tumor supernatant and some nutrient depletion conditions, these data support a model in which acute loss in mTOR signaling and/or translational capacity dampens acute immune function, while prolonged ISR engagement enforces adaptive programming supporting metabolic and functional recovery in nutrient stressed CD8+ T cells.

To understand how changes in mTORC1 signaling, ISR signaling, and global translation contribute to the selective regulatory responses we observed early in the response to nutrient stress, we performed RNA-seq and polysome profiling on cells treated with halofuginone, Torin1, or 4EGi-1 for 3 hours. Whereas halofuginone only mildly impacted polysome abundance, Torin and 4EGi-1 treatment showed a strong decrease in polysome abundance and an increase in monosome abundance, in concordance with their known activities as inhibitors of global translation (Fig 3G). We observed that halofuginone induced transcripts known to be associated with the ISR, including genes involved in amino acid transport, amino acid synthesis, and tRNA charging, just as we observed in our analysis of nutrient-stressed cells (Fig 3H). On the other hand, cells treated with Torin and 4EGi-1 showed a relative depletion of transcripts associated with the cytosolic ribosome, mitochondrial ribosome, and mitochondrial respiratory chain in the polysome-associated RNA pool relative to input RNA, suggestive of translational repression (Fig 3H). In keeping with this, we found that Torin or 4EGi-1 treatment led to a rapid decrease in mitochondrial oxygen consumption rate, while halofuginone treatment led to only a mild decrease in oxygen consumption rate. (Fig 3I). On the other hand, ISR activation drove increased expression of transcripts related to amino acid transport and synthesis including *Slc7a11, Slc6a9, Slc1a4, Slc38a2,* and *Gpt2*, reminiscent of the signature we observed in CD8+ T cells under nutrient stress (Fig S5G). Together, these data suggest that diminished mTOR signaling and/or dampening of cap-dependent translation is responsible for the repression of cellular biosynthesis and growth-related programming we observed during nutrient stress, while the ISR induction drove adaptive metabolic reprogramming and amino acid handling that supports the resolution of nutrient stress and functional recovery.

Lastly, given that rescue of immune function under nutrient stress was associated with restoration of amino acid levels, we next tested if ISR induction, loss of mTOR signaling, or dampening of 4EGi-sensitive translation impacted the amino acid profile of CD8+ T cells. We observed that Torin and 4EGi-1 treatment both led to broadly elevated intracellular amino acid levels, consistent with a decrease in translation-driven amino acid demand, while halofuginone drove a more selective change in the cellular amino acid profile – including serine, alanine, asparagine, proline, and glutamate (Fig 3J). Overall, these data indicate that diminished translational demand leads to a broad elevation of amino acid availability, while ISR-induced gene expression drives more selective changes in amino acid abundance without compromising translational capacity.

Altogether, these findings are consistent with a model where the loss of mTOR signaling and cap-dependent translation in conjunction with ISR activation drives complementary metabolic and regulatory responses that shape functional capacity during T cell nutrient stress adaptation (Fig 3K). Short-term loss of mTOR signaling and translational capacity reduces global biosynthetic and mitochondrial activity, both limiting immune function and reducing demand for amino acids. Paralleling this, ISR engagement drives a more selective metabolic response through the induction of amino acid transport and synthesis transcripts, allowing for long-term functional recovery without wholesale loss of translational or mitochondrial capacity.

### The ISR-driven transcription factors ATF4 and CEBPG are required to support immune function under stress

As activation of the ISR was sufficient to drive elevated function, we next tested if ISR-driven responses were indeed necessary to support CD8+ T cells under nutrient stress (Fig S6A). To do this, we first utilized an inhibitor of GCN2-driven ISR signaling, GCN2iB^61^. GCN2 inhibition impaired the induction of ATF4 downstream of halofuginone treatment and decreased cell viability and CD98 expression after BCAA, arginine, or methionine depletion, but not glucose depletion or control conditions, consistent with the amino acid-related sensing activity of GCN2 (Fig S6B and C). We further observed that GCN2iB-treated CD8+ T cells displayed an inability to adapt to halofuginone treatment. These cells exhibited an inability to enhance cytokine production in response to ISR engagement and instead underwent cell death, in contrast to cells treated with GCN2iB alone (Fig S6D). These results highlight the requirement for ISR-induced programming to sustain cell viability and function under GCN2-activating stresses.

The ISR exerts its effects both translationally via eIF2α phosphorylation and transcriptionally through transcription factors (Fig S6A).^7–9^ While ATF4 is the most well characterized ISR transcription factor, recent reports have suggested that other basic leucine zipper (bZIP) transcription factors, such as C/EBP family members, can dimerize with ATF4 to modify its activity and modulate the outcome of ISR signaling.^56,62–64^. As we observed the activation of a broad network of transcription factors upon acute stress as well as motifs for ATF4 and CEBPG near many of the gene loci whose transcripts were strongly induced during stress, we tested whether ATF4 and/or CEBPG were required to support effector function in stressed CD8+ T cells. To do this, we utilized CRISPR/Cas9 to target Atf4, Cebpg, or the Rosa26 locus as a control, and then evaluated CD8+ T cell function in the presence or absence of GCN2-mediated stress by halofuginone (Fig 4A). By employing electroporation after initial T cell activation, we sought to bypass any confounding role for these factors during initial activation, focusing on their role during the effector phase of the response. We found both ATF4 and CEBPG were selectively required to sustain maximal cytokine production in the presence of halofuginone, while the absence of these factors had little impact on cytokine production in control conditions (Fig 4B). Moreover, loss of either ATF4 or CEBPG dampened the stress-induced increase in CD98, highlighting a shared role for both factors in mediating adaptive metabolic responses under stress (Fig 4B). Given both ATF4 and CEBPG supported T cell function selectively under environmental stress in vitro, we next asked whether these transcription factors were similarly essential for anti-tumor T cell responses in vivo. To test this, we adoptively transferred ATF4- or CEBPG-knockout, P14 TCR-transgenic CD8+ T cells into mice bearing subcutaneous gp33-expressing MC38 tumors and measured tumor growth over time (Fig 4A). In contrast to control cells, both ATF4 and CEBPG knockout CD8+ T cells failed to suppress tumor growth, showing no improvement over animals that did not receive T cells (Fig 4C). Altogether, this suggests that the ISR drives ATF4 and CEBPG expression, which in turn maintain immune function under environmental stress and are required for optimal immunity against solid tumors.

**Figure 4:**
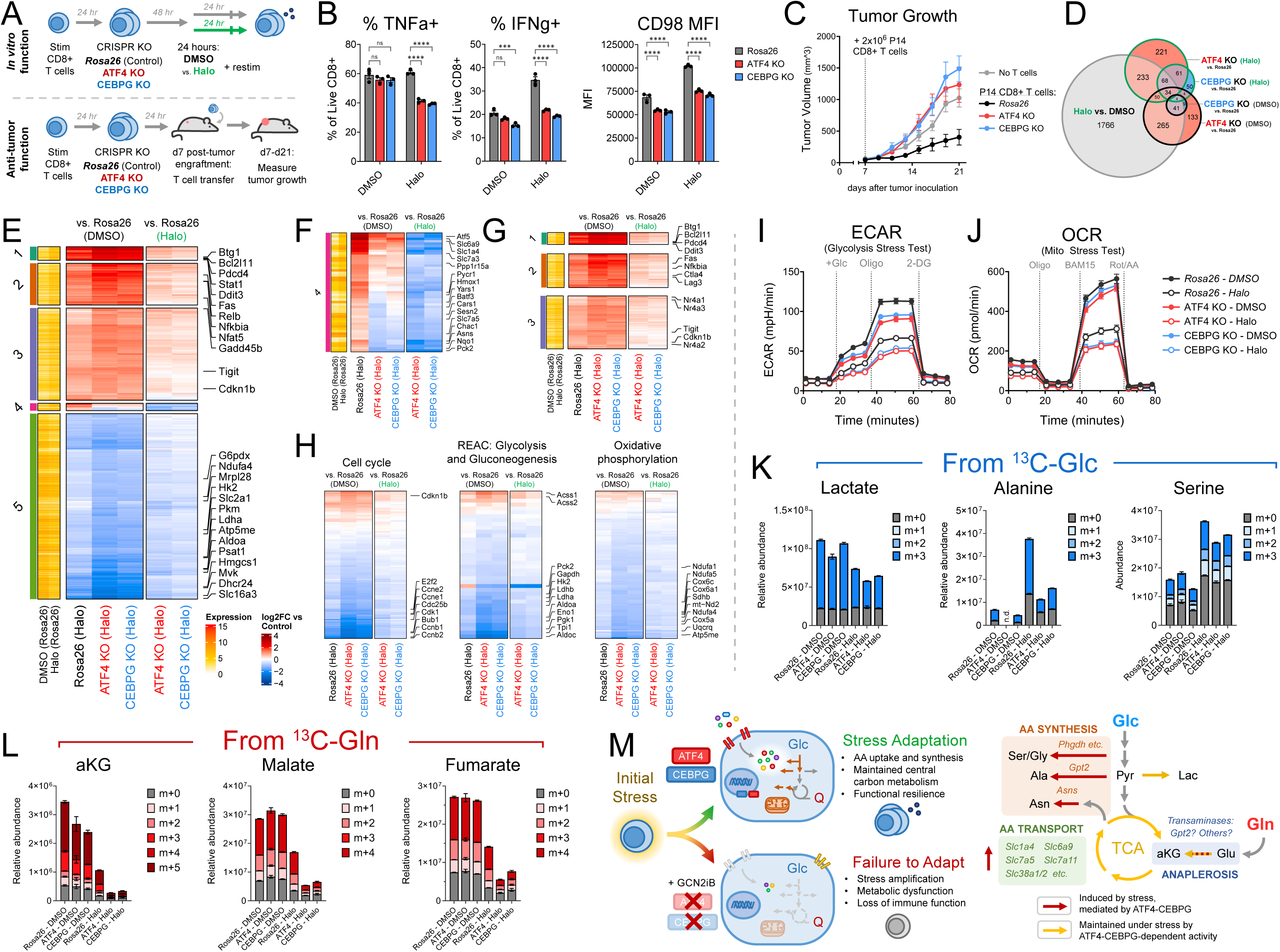
A GCN2-ATF4-CEBPG axis drives metabolic rewiring to prevent runaway stress-driven dysfunction and maintain immune function under stress. A) Schematic of experimental approach for assessing the contribution of ATF4 / CEBPG-driven programming on immune function under stress. Top: *Rosa26* (control), ATF4 KO or CEBPG KO CD8+ T cells were cultured in the presence or absence of halofuginone for 24 hours and restimulated during the final 6 hours of culture. Bottom: *Rosa26* (control), ATF4 KO or CEBPG KO P14 CD8+ T cells were transferred into gp33-MC38 tumor-bearing mice to assess anti-tumor function. B) Intracellular cytokine production and CD98 expression in *Rosa26* (control), ATF4 KO, or CEBPG KO CD8+ T cells in the presence of DMSO or halofuginone for 24 hours. C) gp33-MC38 tumor growth in recipient mice receiving an adoptive transfer of 2×10^6 *Rosa26* (control), ATF4 KO, or CEBPG KO P14 CD8+ T cells at day 7. D) Euler diagram of differentially expressed genes due to ATF4 (red fill) / CEBPG (blue fill) KO in the presence of DMSO (black outline) or halofuginone (green outline); or due to halofuginone treatment (grey) E) Heatmap of all differentially-expressed transcripts in *Rosa26*, ATF4 KO, and CEBPG KO CD8+ T cells cultured in the presence or absence of halofuginone for 24 hours. Left annotation: k-means cluster membership and expression value in *Rosa26* control condition. Left main: Data from halofuginone-treated conditions are plotted relative to *Rosa26* cells cultured in DMSO (data represents the combined effect of halofuginone treatment and gene knockout). Right main: Data from halofuginone-treated conditions are plotted relative to *Rosa26* cells treated with halofuginone (data represents relative change in gene expression attributed to gene KO alone while under stress). F) Heatmap of changes in transcript expression for genes in Cluster 4. G) Heatmap of changes in transcript expression for genes in Clusters 1-3. H) Heatmaps of changes in transcript expression for all annotated genes from indicated pathways. I-J) Oxygen consumption rate (OCR) (I) and extracellular acidification rate (ECAR) (J) in *Rosa26* (control), ATF4 KO, or CEBPG KO CD8+ T cells in the presence of DMSO (closed) or halofuginone (open) for 24 hours. K) Labeling of selected metabolites from U-13C glucose over final 18 hours in *Rosa26* (control), ATF4 KO, or CEBPG KO CD8+ T cells in the presence of DMSO or halofuginone for 24 hours. L) Labeling of selected metabolites from U-13C glutamine over final 18 hours in *Rosa26* (control), ATF4 KO, or CEBPG KO CD8+ T cells cultured in the presence of DMSO or halofuginone for 24 hours. M) Models for role of ATF4 and CEBPG in mediating metabolic adaptation and immune function in CD8+ T cells under stress

### ATF4 and CEBPG coordinate adaptive metabolic rewiring to restrain runaway stress

To understand how ATF4 and CEBPG sustain CD8+ T cell function under environmental stress, we performed RNA-seq on control (*Rosa26)*, ATF4 KO, or CEBPG KO T cells in the presence or absence of halofuginone (Fig 4D and E. We found that long-term ISR induction led to widespread transcriptional reprogramming, including many of the metabolic genes previously observed to be elevated upon acute nutrient stress, such as amino acid transporters and synthesis enzymes like *Slc7a5, Asns,* and *Pycr1.* (Fig 4E and F, Fig S7A). However, we also observed that long-term halofuginone treatment drove the expression of a dysfunction-related gene signature, including increased expression of anti-proliferative programming (*Btg1, Cdkn1b)*, proapoptotic programming (*Fas, Ddit3*), and inhibitory receptors (*Tigit, Lag3*), along with a widespread decrease in expression of transcripts related to cell cycle progression, central carbon metabolism, and cholesterol biosynthesis (Fig 4E-H, Fig S7A and B). While ATF4 and CEBPG were required to support the expression of a discrete module of largely amino-acid-metabolism-related genes (Cluster 4), we noted that loss of these factors led to an overall amplification of dysfunction-related programming (Fig 4E and F, Fig S7A). Concordantly, we observed increased p-eIF2a in ATF4 KO cells and increased ATF4 levels in CEBPG KO cells treated with halofuginone, suggesting dysregulation of the stress response (Fig S7C). These data illustrate a distinction between adaptive and maladaptive programming within the broader transcriptional response to stress. Whereas environmental stress primes gene expression related to both cell dysfunction and cell adaptation, ATF4/CEBPG engage a selective sub-network of targets that mitigate stress amplification and cell dysfunction.

While ATF4 and CEBPG KO cells showed similar profiles of differentially-expressed genes under stress, the overlap in differentially-expressed genes between control and stress conditions for each individual transcription factor was much lower, indicating that these factors have highly context-specific activities (Fig 4D). ATF4 or CEBPG KO also impacted the stress-induced expression of a variety of other transcription factors, including *Atf5, Batf3,* and *Nr4a1,* highlighting their role in supporting a larger transcription factor network (Fig S7D).

Given loss of ATF4 and CEBPG led to widespread changes in metabolic gene expression, we next sought to test their roles in supporting metabolic function in CD8+ T cells under stress. We observed that long-term halofuginone treatment resulted in a decrease in both the glycolytic and respiratory rate of CD8+ T cells (Fig 4I and J, Fig S8A). Here, loss of either ATF4 or CEBPG further decreased rates of glycolysis and mitochondrial respiration during stress, compared to *Rosa26* control cells (Fig 4I and J, Fig S8A). We further found that loss of ATF4 or CEBPG impaired glycolysis in control conditions, indicating a broader role in supporting T cell glycolytic capacity. (Fig 4I, Fig S8A). Altogether, these data indicate that while an ongoing stress response dampens central carbon metabolism, ATF4 and CEBPG support the maintenance of glycolysis and oxidative phosphorylation during stress.

Lastly, we aimed to understand the precise metabolic functions controlled by ATF4 and CEBPG that allow for T cells to maintain function in the face of ongoing stress. To do so, we performed stable isotope labeling using U-^13^C-labeled glucose or glutamine in ATF4 and CEBPG knockout T cells, both in the presence or absence of halofuginone. Consistent with our Seahorse results, we observed that halofuginone drove decreased glucose-derived lactate production, an effect that was further diminished with ATF4 or CEBPG KO (Fig 4K, Fig S8B). ATF4 and CEBPG KO cells treated with halofuginone also displayed decreased abundance of glycolytic intermediates (Fig S8B). Conversely, we observed that halofuginone drove increased production of glucose-derived amino acids – including alanine, serine, and glycine – in an ATF4- and CEBPG-dependent manner (Fig 4K, Fig S8C). As alanine and serine/glycine are known to be important for T cell expansion and function, the observations that serine/glycine/alanine levels and synthesis enzyme expression are both stress-inducible and ATF4/CEBPG-dependent suggest that these pathways contribute to functional resilience in effector CD8+ T cells^65–68^.

Corroborating our Seahorse results, we also observed that halofuginone led to a reduction in both glutamine- and glucose-derived carbons entering the tricarboxylic acid (TCA) cycle (Fig 4L, Fig S8D). Here, ATF4 and CEBPG were selectively required to sustain residual TCA cycle activity, with further losses of TCA cycle metabolite abundance and glutamine-derived labeling of TCA cycle metabolites in ATF4 and CEBPG knockout cells treated with halofuginone (Fig 4L, Fig S8D and E). As the transaminase activity of some amino acid synthesis enzymes like Gpt2 facilitate the production of alpha-ketoglutarate, these data indicate that the ATF4 and CEBPG-driven programming serves to concurrently restore amino acid pools and preserve mitochondrial function in CD8+ T cells under stress. At the same time, we observed a buildup in aspartate levels in ATF4 and CEBPG knockout cells treated with halofuginone, suggestive of dysregulated aspartate-asparagine homeostasis resulting from decreased expression of the ATF4 / CEBPG target asparagine synthetase (Asns) (Fig 4F, Fig S8D and E). Lastly, we found that levels of the branched chain amino acids leucine and isoleucine were decreased in ATF4 and CEBPg KO cells treated with halofuginone, consistent with our observation that both factor are required to support stress-driven increases in CD98 expression (Fig 4B, Fig S8F). These data suggest that ATF4 and CEBPG support central carbon metabolism, and amino acid synthesis, and amino acid uptake under stress.

Collectively, our findings support a model whereby ATF4 and CEBPG orchestrate functional resilience of CD8+ T cells under environmental stress through an “adaptive” metabolic program supporting amino acid accumulation and preservation of central carbon metabolism (Fig 4M). Here, ATF4- and CEBPG-dependent regulation of metabolic gene expression (including increases in amino acid synthesis enzymes, amino acid transporters, and transaminases such as *Gpt2*) mitigates the maladaptive effects of environmental stress and preserves protective immune function despite dampened central carbon metabolism. In the absence of these factors, CD8+ T cells under stress fail to maintain amino acid levels and mitochondrial anaplerosis, leading to collapse of central carbon metabolism, amplification of a dysfunction-related transcriptional profile, and compromised immune function.

## Discussion

Gene-regulatory mechanisms that control metabolism and cell behavior in response to environmental perturbation have been a longstanding focus of cell and molecular biology^69–71^. Adaptive metabolic responses to extracellular nutrient levels are observed even among unicellular organisms, allowing cellular life to grow or persist across diverse biochemical conditions^4,10,72–76^. In multicellular organisms, however, individual cells must perform lineage-specific functions that promote the overall health of the organism, balancing the use of energy and nutrient stores for both functional programs and primordial cellular processes such as growth and self-maintenance. Studies of transformed cells have helped reveal how both intrinsic and environmental signals regulate cell growth and metabolism in multicellular systems, as well as how cells are able to grow or survive — in spite of environmental stress and at the expense of the multicellular host — when these regulatory systems fail^1,2,77,78^. Despite this, how non-transformed cell lineages prioritize nutrient use across programs of growth, survival, and/or lineage-specific functions while under nutrient limitation or environmental stress remains comparatively unknown. Additionally, factors that allow non-transformed cells to maintain key functions despite the suboptimal environmental conditions that may arise during disease may prove to be important targets for therapeutic application. As effector T cells are tasked with providing protective immunity in metabolically-disrupted sites such as infected tissue or tumors, these cells offer a dynamic system to better understand the coordination of metabolism, gene expression, and nutrient allocation under environmental stress with the potential for novel therapeutic application in immunotherapy.

We propose that “biosynthetic plasticity” — the ability to rapidly alter global nutrient handling and reprioritize nutrient allocation across discrete metabolic and cellular programs in response to environmental change — represents a novel mechanism preserving lineage-specific functionality in a wider array of biochemical contexts. In comparison to plasticity in fuel choice, in which cells readily utilize varying carbon sources to maintain energetic capacity resulting in distinct functional outcomes, biosynthetic plasticity allows cells to reprioritize metabolic networks to sustain an existing functional program at a diminished energetic capacity^33,66,79–88^. In the case of our work in CD8+ T cells, we find that cells under acute stress rapidly modulate nutrient-sensitive signaling and global translational activity, reconfiguring cellular nutrient handling while shifting the translational priority of discrete cellular pathways. This early response licenses an adaptive transcriptional program to inducibly establish an alternate mode of metabolism that mitigates cell dysfunction and metabolic collapse under environmental stress, promoting functional resilience and potentiating anti-tumor immunity.

Our work highlights the complex interplay between pro-growth signaling, stress signaling, and translation in T cells, adding to a growing body of literature evincing a key role for translational control and metabolism-translation cross-talk in T cell responses^30,31,48,103–113^. We find the mTOR and ISR pathways shape immune function under stress by jointly governing metabolic activity, translational capacity, and regulatory programming, consistent with similar findings in other cellular systems^10,50,56,58,59,89–93^. In T cells, mTORC1 activity is known to support T cell effector differentiation, proliferation, and function while stress-responsive signaling has been shown to exert both adaptive and maladaptive outcomes in T cells^7–9,34,37,94–102^. Indeed, we observed that loss of immune function under acute nutrient stress coincides with decreased mTORC1 signaling and increased ISR signaling. Additionally, we observed loss of mTORC1 activity over longer periods dampens T cell function. Despite this, we observed that loss of mTORC1 signaling decreases global translational activity, increasing intracellular amino acid content and repressing the translation of growth-related programming. Concurrently, the ISR induces an adaptive program of amino acid transporters, synthesis enzymes, and transcription factor transcripts that is efficiently translated under stress and correlates with rescued or enhanced immune function over time. Thus, we propose that these signaling responses act cooperatively to harmonize global translational activity and biosynthetic activity with the constraints of the metabolic environment, promoting cellular resilience to nutrient stress and allowing for more robust protective immunity across varying metabolic microenvironments.

In addition to mRNA translation-driven mechanisms, we highlight a key module within a larger stress-sensitive transcriptional network that supports long-term stress resilience and immune function. In the absence of ATF4 or CEBPG, CD8+ T cells show dampened immune function under stress, dysregulated central carbon metabolism, amplification of dysfunction-related programming, and compromised anti-tumor efficacy. This occurs even though ATF4 and CEBPG directly regulate a narrow subset of stress-sensitive genes (largely composed of amino acid transport and synthesis enzymes), highlighting the importance of ATF4 / CEBPG-dependent metabolic programming in preserving T cell function by averting dysfunctional fates under stress. In keeping with increasing evidence of metabolic contributions to T cell dysfunction and exhaustion, we observe that ATF4 and CEBPG KOs under stress show increased expression of dysfunction-related targets (e.g. Btg1, Nr4a1, Lag3, Tigit)^23–26,114–122^. Additional work is required to dissect how environmental stress and stress-sensitive transcriptional programs contribute to the variety of dysfunctional CD8+ T cell states. Of note, we observed that while the regulomes of ATF4 and CEBPG under stress are quite similar, each individual factor’s regulome differs significantly between stress and control conditions, suggesting that the environmental context shapes the regulome of individual transcription factors. This observation, as well as the finding that ATF4 / CEBPG KO alters expression of multiple other transcription factors in cells under stress, highlights the dynamic and interconnected nature of stress adaptation responses over time and presents an opportunity for further dissection of environmental feedback on stress-sensitive regulatory and metabolic networks. As these factors are amongst dozens we observed to be acutely nutrient-sensitive, there are likely additional regulators of stress resilience to be identified.

Beyond this study, there remains much to learn about how specificity in stress-induced regulatory responses is achieved, particularly within lymphocytes where these phenomena are underexplored. It will be important to understand the precise molecular mechanisms through which this regulation is enforced, including alternate modes of translation, upstream open reading frames, unique RNA sequence elements, post-transcriptional modifications, and direct control of translation through regulatory RNAs or RNA-binding proteins^44,45,47,103,104,112,113,123–132^. Manipulation of these mechanisms would open avenues to preserving translational “priority” to enforce expression of desired transcripts for cell therapies. Additionally, new screening modalities may allow for a systematic dissection of the combinatorial TF-TF and TF-environment interactions that underlie environmental stress responses as well as the diversity of stress-adapted and dysfunctional cellular states in T cells and other non-transformed cell types^133–135^. While our study focuses on CD8+ T cells, it will be important to understand how environmental stressors interact with the functional programming of multiple immune cell lineages within complex microenvironments to broadly shape immune responses.^136–141^ This will enhance our understanding of stress resilience in immunity to add to the growing number of strategies utilizing metabolic conditioning or cell engineering for therapeutic applications^20,22,102,142–144^ Additionally, as sites of peripheral immune function may contain multiple cell types and several overlapping stresses in addition to nutrient limitation (e.g. inflammation, hypoxia, or ongoing therapy), additional models of immune cell adaptation across microenvironmental, tissue, and disease contexts will allow for a more complete understanding of the impact of environmental stress on immunity^136,145,146^. Lastly, the stress-sensitive targets across our datasets, including those with little annotated function, may prove useful for the discovery of conserved factors that promote stress-resilience or susceptibility to be therapeutically targeted beyond the immune system - for example, in cancer, where stress adaptation pathways, translational and metabolic status, and environmental conditions modulate therapeutic efficacy or resistance^60,147–162^.

## Methods

### Mice

C57BL/6J (B6, Jax #000664), Tg(TcraTcrb)1100Mjb (OT-I, Jax #003831), B6.Cg-Tcratm1MomTg(TcrLCMV)327Sdz/TacMmjax (P14, Jax #037394), and B6.SJL-Ptprca Pepcb/BoyJ (CD45.1, Jax #002014) mice were obtained from The Jackson Laboratory and maintained under specific pathogen-free conditions. Experiments were performed with male or female mice aged between 6 and 9 weeks of age. All experiments were performed in accordance with the Institutional Animal Care and Use Committee of the Children’s Hospital of Philadelphia (IAC 21-001325).

### CD8+ T cell Isolation and Stimulation

Bulk mouse CD8+ T cells were isolated from spleen and peripheral lymph nodes by ACK-lysis and magnetic negative selection using an EasySep™ Mouse CD8+ T Cell Isolation Kit (STEMCELL Technologies, #19853). For experiments without electroporation, purified CD8+ T cells were initially stimulated for 24 hours on tissue culture plates coated with 2.5ug/mL anti-CD3ε (clone 145-2C11, BioLegend, #100360, RRID:AB_2800555) and 2.5u/.mL anti-CD28 antibodies (clone 37.51, BioLegend, #102122, RRID:AB_11147170), then removed from stimulation for further culture. For experiments involving electroporation, cells were stimulated for 24 hours on plates coated with 5ug/mL anti-CD3ε and 5ug/mL anti-CD28 antibodies before electroporation.

### Cell Lines

MC38 cells were a gift from the lab of Ken Cadwell (University of Pennsylvania, Philadelphia, PA). gp33-expressing MC38 cells were a gift from the lab of E. John Wherry (University of Pennsylvania, Philadelphia, PA). MC38 and gp33-MC38 cells were maintained at <80% confluence by trypsinization and passaging every two days for one week before use in further experiments. Cells were cultured in Dulbecco’s Modified Eagle Medium (DMEM, Gibco #11965118) 10% fetal bovine serum Corning, #MT35071CV), additional 2mM L-glutamine (Gibco, #25030081), and 1x penicillin/streptomycin (Gibco, #15140122).

### Media and Cell Culture

Initial T cell stimulation and culture was performed in supplemented lymphocyte culture medium (“T cell culture medium”, TCM): Roswell Park Memorial Institute (RPMI) 1640 Medium (Gibco, #11875093), supplemented with 10% fetal bovine serum (Corning, #MT35071CV), additional 2mM L-glutamine (Gibco, #25030081), 1mM sodium pyruvate (Gibco, #11360070), 25mM HEPES (Gibco, #15630080), 1x penicillin/streptomycin (Gibco, #15140122), 55uM 2-mercaptoethanol (Gibco, #21985023), and 5ng/mL recombinant mouse IL-2 (BioLegend, #575406).

MC38-conditioned supernatant was generated by passaging MC38 cells for one weekend allowing the cell cultures to reach 80% confluence, then replacing the culture media with T cell culture media (above) for 48 hours. Supernatant was then centrifuged for 15m at 4000xg to remove cells and utilized for further experiments.

All nutrient-stress media contained an RPMI-derived base medium: “Control” = RPMI-1640 Medium (Gibco, #11875093); No Glucose = RPMI-1640 Medium, no glucose (Gibco, #11879020); No Glutamine = RPMI 1640 Medium, no glutamine (Gibco, #21870076); No BCAA, RPMI 1640 Medium w/ L-Glutamine, w/o L-Isoleucine, L-Leucine, L-Valine (US Biological, #R8999-20); No Arginine = RPMI 1640 Medium for SILAC (Gibco, #A33823) + 40mg/L-Lysine (Thermo, #J62225.36); no Methionine = RPMI 1640 Medium w/o L-Glutamine, Methionine (US Biological, #R8999-06) + 2mM L-Glutamine (Gibco, #25030081)), 10% dialyzed fetal bovine serum (Cytiva, #SH30079.03), additional 2mM L-glutamine (except “No Glutamine” condition, Gibco #25030081), 1mM sodium pyruvate (Gibco, #11360070), 25mM HEPES (Gibco, #15630080), 1x penicillin/streptomycin (Gibco, #15140122), 55uM 2-mercaptoethanol (Gibco, #21985023), and 5ng/mL recombinant mouse IL-2 (BioLegend, #575406).

All cells were cultured in a CO2 incubator at 37C at 5% CO2.

### Pharmacological Agents

All experiments utilizing pharmacological agents below were performed in “Control” media as described in the nutrient-stress media section above. All compounds were resuspended in DMSO (VWR #25-950-CQC) and used at the following final concentrations unless otherwise specified: Torin 1 (500nM, Cayman Chemical, #10997), Halofuginone (hydrochloride) (50nM, Cayman Chemical, #13370), 4EGi-1 (50uM, Cayman Chemical, #15362), GCN2iB (2uM, Cayman Chemical, #35897).

### CRISPR-Cas9 KO by Electroporation

KO cells were generated via electroporation of Cas9-RNPs using a Thermo Neon instrument. Cas9-RNPs were generated by combining by 1.25uL 200uM target-specific crRNA (IDT), 1.25uL 200uM ATTO-550-labeled tracrRNA (IDT, #1075928), and 2.5uL nuclease-free duplex buffer (IDT) and incubating for 5m at 95C, then mixed with 5uL Cas9-NLS (QB3-Berkeley MacroLab) and incubated for 15m before use in electroporation. 2-10 x 10^6^ activated T cells were washed once with PBS, resuspended in 100uL Buffer R (Thermo #MPK10096), mixed with 10uL Cas9-RNP, then electroporated with the following parameters (1600V, 10ms, 3 pulses) and cultured in “T Cell Media” without Pen/Strep for 24 hours before further use. Viability and RNP uptake were quantified at 24 hours post-nucleofection by flow cytometry.

crRNA target sequences:

Mouse *Rosa26 locus*: ACTCCAGTCTTTCTAGAAGA
Mouse Atf4: AAGTTTAATAAAAGTCGACC
Mouse Cebpg: CAGAGAGCGGAACAATATGG

### Restimulation and Flow Cytometry

All flow cytometry was performed on a CytoFLEX LX or CytoFLEX S cytometer (Beckman Coulter). Intracellular cytokine production was measured by adding brefeldin A (Invivogen) 30 min after transferring cells to a plate coated with 2.5ug/mL anti-CD3ε (clone 145-2C11, BioLegend, #100360, RRID:AB_2800555) and 2.5ug.mL anti-CD28 antibodies (clone 37.51, BioLegend, #102122, RRID:AB_11147170) and culturing for 6 hours before staining. Cells were first stained with surface antibodies and viability dye (30m at RT) diluted in PBS, fixed with 1x Fixation/Permeabilization Buffer for 15m at room temperature (eBioscience™ Foxp3 / Transcription Factor Staining Buffer Set, Invitrogen, #00-5523-00), and lastly stained with intracellular antibodies diluted in 1x Permeabilization Buffer (eBioscience™ Foxp3 / Transcription Factor Staining Buffer Set, Invitrogen, #00-5523-00) for 30m at room temperature. Reagents utilized: eBioscience Fixable Viability Dye eFluor 780 (1:1000, Thermo Fisher Scientific, #65-0865-18); CD8 alpha Monoclonal Antibody (53-6.7), FITC (1:300, Invitrogen #MA1-10303, RRID: AB_11153636); CD98 Monoclonal Antibody (RL388), PE, eBioscience™ (1:300, Invitrogen #12-0981-81, RRID:AB_465792); Granzyme B Monoclonal Antibody (NGZB), PE-eFluor™ 610, eBioscience™, (1:300, Invitrogen, #61-8898-82, RRID: AB_2574670); IFN gamma Monoclonal Antibody (XMG1.2), APC, eBioscience™ (1:300, #17-7311-82, RRID: AB_469504); TNF alpha Monoclonal Antibody (MP6-XT22), eFluor™ 450, eBioscience™ (1:300, #48-7321-82, RRID:AB_1548825)

### RT-qPCR

RNA was isolated with the Zymo Quick-RNA MicroPrep Kit (Zymo Research, #R1050) with DNase I treatment according to the manufacturer’s instructions. cDNA was synthesized with 1ug of purified RNA using Thermo Maxima H Minus Reverse Transcriptase (Thermo Scientific, #EP0751) according to manufacturer’s instructions using 25 pmol oligo(dT)_18_ (Thermo Scientific, #SO132) and 25 pmol random hexamer (Thermo Scientific, #SO142) primers and the following thermocycler protocol: 10 min at 25°C, 15 min at 50°C, 5 min at 85°C. cDNA was diluted 1:10 in molecular biology water and quantified via real-time quantitative polymerase chain reaction using PowerTrack™ SYBR Green Master Mix (Thermo Scientific, #A46110) and 1.66uM forward and reverse primers on a Bio-Rad CFX96 or CFX384 instrument (Bio-Rad). 3-4 technical replicates were averaged, each quantified transcript was normalized to *Rps18*, and data from all samples were plotted as relative expression versus a chosen sample.

Mouse Atf3 F – AGAGTGCCTGCAGAAAGAGTCA
Mouse Atf3 R – CGGTGCAGGTTGAGCATGTAT
Mouse Ppp1r15a F – TCCTCTAAAAGCTCGGAAGGTACAC
Mouse Ppp1r15a R – TCTCGTGCAAACTGCTCCCA
Mouse Sesn2 F – ACCTTCCGTGCCCAGGATTAT
Mouse Sesn2 R – CCTGGAACTTCTCATCCAGCAG
Mouse Nupr1 F – CTGGCGGGCATGAGAGGAAG
Mouse Nupr1 F – TCTGTGGTCTGGCCTTATCTCCA
Mouse Slc3a2 F – CACTGGGGAGCGTACTGAATCC
Mouse Slc3a2 F – AGTCCTGCTTGCGACACACTC
Mouse Slc7a5 F – AGCTGTGGCTGTGGACTTCG
Mouse Slc7a5 R – CCCATTGACAGAGCCGAAGCA
Mouse Slc7a11 F – CCAAGGGCATACTCCAGAACA
Mouse Slc7a11 R – TAGGACAGGGCTCCAAAAAGT

### Western Blot

Cells were pelleted, washed in PBS and lysed in RIPA lysis buffer (50 mM Tris (pH 8.0), 1 mM EDTA, 150 mM NaCl, 1% NP-40, 0.5% Na-deoxycholate, 0.1% SDS) containing 10 mM NaF, 1 mM Na3VO4, and cOmplete™, Mini, EDTA-free Protease Inhibitor Cocktail (Roche, #11836170001). Protein lysate concentration was quantified via Bradford assay (Bio-Rad, #5000006) using bovine serum albumin as standard. 15-20ug protein lysate was mixed with reducing sample buffer (Boston BioProducts, #BP-111R), and samples were separated by SDS-PAGE on Any kD™ Mini-PROTEAN® TGX™ Precast Protein Gels (BioRad, #456-8126) followed by transfer to a nitrocellulose membrane using the Trans-Blot Turbo Transfer System (BioRad) on the Turbo setting. Membranes were blocked with 5% w/v dry nonfat milk in Tris-buffered saline containing 0.1% Tween 20 (TBS-T) for 60 min. Primary antibodies were diluted in 1x Tris Buffered Saline (TBS) with 1% Casein (BioRad, #1610782) overnight at 4°C with shaking. Five TBS-T washes of 2 minutes each were performed before incubation with secondary antibody (diluted in 5% w/v dry nonfat milk in TBS-T) for 2 hours at room temperature with rocking. Three additional washes were performed before visualizing the chemiluminescent signal with SuperSignal West Pico PLUS (Thermo Scientific, #34580) or Femto Chemiluminescent Substrate (Thermo Scientific, #34095) on a ChemiDoc MP Imaging System (BioRad). For phospho-epitopes, blots were stripped using Restore™ Western Blot Stripping Buffer (Thermo Scientific, # 21059) after visualization of phospho-epitopes before incubating with total antibody. Actin was quantified by incubation with anti-actin fAb at room temperature for 1h. For western blot time courses involving multiple gels, an identical (pooled) protein sample from cells at t=0 of the experiment was loaded on every gel as an internal control to account for changes in transfer efficiency. All membranes within the time course were imaged for the same amount of time for a given target.

Phospho-S6 Ribosomal Protein (Ser235/236) (D57.2.2E) XP® Rabbit mAb, 1:5000 (Cell Signaling Technology, #4858)

S6 Ribosomal Protein (5G10) Rabbit mAb, 1:5000 (Cell Signaling Technology, #2217)

ATF-4 (D4B8) Rabbit mAb, 1:1000 (Cell Signaling Technology, #11815)

Goat anti-Rabbit IgG (H+L) Cross-Adsorbed Secondary Antibody, HRP (Invitrogen #G-21234, RRID: AB_2536530)

Goat anti-Mouse IgG (H+L) Secondary Antibody, HRP (Invitrogen, #A16066, RRID:AB_2534739) hFAB™ Rhodamine Anti-Actin Primary Antibody, 1:2000 (Bio-Rad, #12004164)

### Polysome Fractionation

Sucrose solutions were prepared in polysome extraction buffer (10 mM HEPES, 100 mM KCl, 5 mM MgCl2, pH 7.4, 100 ug/ml Cycloheximide, 2 mM DTT). Sucrose gradients were prepared in SW41 ultracentrifuge tubes by mixing 15% and 45% sucrose solution using a BioComp Gradient Master 108. 20M cells per sample per condition were resuspended after 3 hours of culture in nutrient stress media, washed once with PBS containing 50ug/mL cycloheximide, and lysed in 500uL polysome extraction buffer containing 1% Triton-X and SUPERAse In (Thermo Scientific, #AM2696), followed by 30min incubation on ice with vortexing at 5min intervals. Lysates were centrifuged for at 12000g for 15 min at 4C, then 50uL lysate was snap frozen as “Input” RNA and 400uL lysate was loaded onto 15–45% sucrose gradients followed by centrifugation at 37K rpm for 2hr at 4°C in a SW41 rotor of an Optima XPN 80 Beckman ultracentrifuge. Gradients were fractionated with a speed of 800uL/min using a Biocomp piston gradient fractionator, which recorded the OD254nm. Fractions corresponding to 60 sec intervals were collected, labeled 1-15, and stored at −80c before RNA extraction.

250uL of each sucrose fraction was aliquoted for RNA isolation. 20uL of input RNA was diluted to 250uL with molecular biology water before processing. 750uL of Trizol LS containing 20pg of luciferase control RNA (Promega, #L4561) was added to each sample and thoroughly mixed. 1mL of 100% ethanol was added to each sample and thoroughly mixed. Each sample was loaded into a separate column of a Zymo Direct-Zol-96 plate and purified according to manufacturer’s instructions. Eluted RNA was quantified on a Nanodrop, and a pooled sample containing an equal volume of the RNA from polysome-bound fractions (fractions 9-13) was generated for downstream sequencing.

### Total RNA-sequencing and mRNA-sequencing

For polysome profiling via total RNA-seq of nutrient stress conditions, RNA samples were sequenced at the Center for Applied Genomics at the Children’s Hospital of Philadelphia. RNA samples were checked for quality using an Agilent tapestation and libraries were prepared using the Illumina TruSeq Stranded Total RNA Gold according to manufacturer’s instructions. Sample input was 300ng, and samples were sequenced on a NovaSeq 6000 utilizing the S2 300 cycle kit v1.5 and demultiplexed using DRAGEN.

For polysome profiling via mRNA-seq of drug conditions, RNA samples were sequenced at Novogene. Samples were checked for quality using a tapestation and libraries were prepped for stranded mRNA sequencing at Novogene. Messenger RNA was purified from total RNA using poly-oligo-attached magnetic beads. After fragmentation, the first strand cDNA was synthesized using random hexamer primers. Then the second strand cDNA was synthesized using dUTP, instead of dTTP. The directional library was ready after end repair, A-tailing, adapter ligation, size selection, amplification, and purification. The library was checked with Qubit and real-time PCR for quantification and bioanalyzer for size distribution detection. After library quality control, different libraries were pooled based on effective concentration and targeted data amount, then subjected to Illumina sequencing.

RNA-Sequencing reads were initially assessed for quality with fastqc (0.12.1)^163^and multiqc (1.14)^164^. Reads were trimmed of any remaining adapters and filtered with Trimmomatic (0.39)^165^ using the following parameters: seedMismatches: 2, palindromeClipThreshold: 30, simpleClipThreshold: 10, Leading: 3, Trailing: 3, Avgqual: 15, SlidingWindow: 4:15, Minlength: 75. Reads passing the quality control checks were aligned to the GRCm39 mouse genome assembly^166^ with GENCODE annotation set vM31^167^ using STAR (2.7.10b)^168^. Samtools (1.16.1)^169^ was used to sort alignment files by coordinate and index. Gene pseudocounts were obtained with htseq-count from HTSeq (0.11.1)^170^.

EdgeR (4.3.0)^171^ was used to filter low expressing genes (filterByExpr), normalize by library size (calcNormExpr), and control for variance between genes (estimateDisp). Principal component analysis (PCA) was performed on scaled logCPM of genes for each sample (obtained using edgeR’s cpm function) with R’s prcomp function. Gene biotypes were pulled using biomaRt (2.61)^172^ to interact with the ensembl database. Polysome-associated RNA-seq results were further compared to total RNA-seq results with the anota2seq R package (1.27.0)^173^ using the following parameters: minSlopeTranslation = −1,maxSlopeTranslation = 2, minSlopeBuffering = −2, maxSlopeBuffering = 1, maxPAdj = 0.05). Sample by sample Euclidean distances were calculated using R’s dist() function and compared between assays with a Wilcoxon test. Plots were generated with either ggplot2 (3.5.1)^174^, UpsetR (1.4.0)^175^, gprofiler2 (0.2.3)^176^, or ComplexHeatmap (2.18.0)^177,178^.

### Seahorse Extracellular Flux Analysis and Resipher

For Seahorse assay, 3 × 10^5^ cells were plated per well in a 96-well Seahorse assay plate precoated with Cell-Tak (Corning, #354240). For the mitochondrial stress test assay, cells were plated in Seahorse XF RPMI medium (Agilent, #103576-100) containing 10mM glucose, 2mM L-glutamine, and 1mM sodium pyruvate and subsequently treated with oligomycin (1.5 μM, Sigma-Aldrich, #O4876), BAM15 (2.5 μM, Cayman Chemical, #17811), and rotenone (0.5 μM, Cayman Chemical, #13995-1) / antimycin A (0.5 μM, Sigma-Aldrich, #A8674) at indicated time points. For the glycolysis stress test assay, cells were plated in glucose-free Seahorse XF RPMI (Agilent, #103576-100) with 2mM L-glutamine and 1mM sodium pyruvate prior to assay and subsequently treated with 10mM glucose, oligomycin (1 μM, Sigma-Aldrich, #O4876), and 2-deoxyglucose (50mM, Sigma-Aldrich, #D0051) at indicated time points. Data transformation was performed using Wave software.

For Resipher assay, 3 × 10^5^ cells per well were plated in a Nunc MicroWell 96 well plate (Thermo Scientific, #167008) and a Resipher sensing lid (Lucid Scientific) was placed on top. 2-4 wells were left empty for baseline correction. After connection with the Resipher hub, sensors were equilibrated for 6 hours before collecting continuous oxygen consumption measurements.

### Adoptive Transfer Tumor Model

Gp33-expressing MC38 cells were passaged for a week in culture before engrafting 200K cells on the right flank of C57BL/6 recipient mice. On days 5, 6, and 7 post-tumor-engraftment, purified CD45.1.2 P14 CD8+ T cells were respectively stimulated, nucleofected with Cas9-RNP complexes against the *Rosa26* locus (control), Atf4, or Cebpg, and 2 x 10^6^ cells were adoptively transferred into anesthetized, tumor-bearing mice via retroorbital injection. Tumor volume was monitored every 2 days using calipers and calculated using the formula (L x W x W) / 2. Experimental groups were randomly assigned and evenly distributed to individual recipients across multiple cages, and the experimenter measuring tumor size was blinded to treatment groups.

### LC-MS Metabolomics

Following cell culture, 1.5 million cells per sample were washed with PBS, pelleted, and flash-frozen. Cell pellets were extracted in 1mL −80C 80% Methanol / 20% Optima H_2_O containing a mix of 13C/15N-labeled amino acid internal standards (Cambridge Isotopes Laboratory, #MSK-A2-1.2). Samples were pulse-sonicated on ice with a sonic dismembranator (Fisher Scientific, Waltham, MA) 30 times over 15 seconds then incubated on ice for 10 min. Debris was pelleted at 12000 x g for 10 min at 4C and the solvent was dried under nitrogen gas using a blowdown evaporator (Organomation, West Berlin, MA). Dry metabolites were re-suspended in 300 μL of 5% MeOH, vortexed for 1 minute, dissolved using an ultrasonic bath for 15 min, spun down at 12000 x g for 10 min at 4C and distributed to HPLC vials for LC-HRMS analysis. A pooled QC sample was generated for monitoring intra-run variance by mixing 20 μL of each re-suspended experimental sample within the run.

Metabolites were separated using a Thermo Hypersil Gold, 150 x 2.1 mm, 1.9 μm using the UltiMate 3000 quaternary UHPLC (Thermo Scientific, Waltham, MA) equipped with a refrigerated autosampler (6C) and column heater (55C). Solvent A was water with 0.1% formic acid and solvent B was methanol with 0.1% formic acid. The gradient was as follows: 0.5% B at 0 min, 0.5% B at 2 min, 50% B at 6 min, 100 % B at 12 min, 100 % B at 16 min, and back to 0.5 % B at 17 min and kept for 3 more min for column re-equilibration. The flow rate was 0.45 ml/min. A Q Exactive HF mass analyzer (QE-HF) (Thermo Scientific, Waltham, MA) equipped with a heated electro-spray ionization (HESI) source was operated in positive mode in full scan at 120,000 resolution, AGC target = 1e6; Maximum IT = 100 ms; scan range, 70 to 800 m/z. The pooled QC samples were used for metabolite identification by generating MS/MS spectra (dd-MS2) of the top 10 features at 15,000 resolution, AGC target = 1e5, Maximum IT = 25 ms, and (N)CE/stepped NCE = 30, 50, 60v.

For data analysis, metabolites were identified based on the exact mass +/-5 ppm and retention time like authentic standards (Sigma Metabolomics Library). Peak integration was conducted using Xcalibur 4.2 (Thermo Fisher Scientific). Normalized metabolite abundances were generated by plotting the ratio of the detected metabolite peak area relative to its corresponding 13C-labeled internal standard or (if not present) the internal standard with the nearest retention time. Data was imported and plotted in R^179^ using the tidyverse family of packages^180^ including ggplot2^174^.

### Stable Isotope Labeling and GC-MS Metabolomics

Following cell culture, 1.5 million cells per sample were washed with PBS, pelleted, and flash-frozen. Cell pellets were extracted in 640uL −80C 80% Methanol / 20% Optima H_2_O. Extracts were dried using a speedvac and then treated with MOX for 1 hour at 60°C followed by a 45-minute incubation with tBDMS at 60°C. Samples were injected on GC-MS (Agilent Technologies) by the Analytical Core for Metabolomics and Nutrition at the British Columbia Children’s Hospital Research Institute. Metabolites were identified with m/z and retention times as shown in Table S1. Mass isotopologue distributions of selected metabolite ion fragments were quantified and corrected for natural isotope abundance using algorithms adapted from Fernandez et al.^181^. The code is available on https://github.com/Sethjparker/IntegrateNetCDF_WithCorrect (Accessed on November 25th. 2024) under MIT license.

### Proteomics

#### Protein Extraction

Pellets were solubilized in 200 µL of extraction buffer containing 5% sodium dodecyl sulfate (SDS, Affymetrix), 50 mM TEAB (pH 8.5, Sigma), and protease inhibitor cocktail (Roche cOmplete, EDTA-free). Each sample (100 µL) was sonicated for 10 minutes at 20°C in a Covaris R230 focused-ultrasonicator (settings: Dithering Y=3.0, Speed=20.0, PIP=360.0, DF=30, CPB=200) to shear DNA and ensure complete solubilization. Samples were then centrifuged at 3000g for 10 minutes to clarify the lysate. Protein concentration was measured by intrinsic tryptophan fluorescence (excitation at 280 nm, emission at 350 nm) against an in-house E. coli lysate standard curve on a Synergy H1 microplate reader (BioTek).

#### In-Solution Digestion

Each sample (130 µg) was digested following the S-Trap (Protifi) manufacturer’s protocol^182^. Briefly, proteins were reduced with 5 mM TCEP (Thermo), alkylated with 20 mM iodoacetamide (Sigma), and acidified with phosphoric acid (Aldrich) to reach a final concentration of 1.2%. Samples were then diluted with 90% methanol (Fisher) in 100 mM TEAB, loaded onto an S-trap column, and washed three times with 90% methanol in 100 mM TEAB. A 1:10 enzyme-to-protein ratio of Trypsin (Promega) and LysC (Wako) in 20 µL of 50 mM TEAB was added, and samples were digested at 37 °C in a humidity chamber for 18 hours. Peptides were eluted sequentially with 50 mM TEAB (40 µL), 0.1% TFA in water (40 µL), and 50/50 acetonitrile:water with 0.1% TFA (40 µL). The combined eluates were dried by vacuum centrifugation, desalted using the Phoenix peptide cleanup kit (PreOmics) per the manufacturer’s protocol, and eluted into autosampler vials. After drying, samples were reconstituted in 0.1% TFA containing iRT peptides (Biognosys, Schlieren, Switzerland). Peptide concentrations were measured at OD280 using a Synergy H1 microplate reader (BioTek) and adjusted to 400 ng/µL for injection.

#### Mass Spectrometry Data Acquisition

Samples were randomized and analyzed on an Exploris 480 mass spectrometer (ThermoFisher Scientific) coupled with an Ultimate 3000 nano UPLC system and EasySpray source. 2ug of each sample was loaded onto an Acclaim PepMap 100 75 µm × 2 cm trap column at 5 µL/min, then separated by reverse phase HPLC on a 75 µm id × 50 cm PepMap RSLC C18 column. Mobile phase A was 0.1% formic acid, and mobile phase B was 0.1% formic acid/acetonitrile. Peptides were eluted into the mass spectrometer at 210 nL/min using a 150-minute gradient from 3% B to 45% B.

Data independent acquisition (DIA) mass spectrometer settings were as follows: one full MS scan at 120,000 resolution, with a scan range of 350-1200 m/z and automatic gain control (AGC) target of 300%, and automatic maximum inject time. This was followed by variable DIA isolation windows, MS2 scans at 30,000 resolution with an AGC target of 1000%, and automatic injection time. The default charge state was 2, the first mass was fixed at 200 m/z, and the normalized collision energy for each window was set at 27.

#### Mass Spectrometry QA/QC and System Suitability

The suitability of the instrumentation was monitored using QuiC software (Biognosys; Schlieren, Switzerland) for the analysis of the spiked-in iRT peptides. As a measure for quality control, standard E. coli protein digest was injected in between samples and data was collected in data dependent acquisition (DDA) mode. The collected data were analyzed in MaxQuant^183^ and the output was subsequently visualized using the PTXQC package^184^ to track the quality of the instrumentation.

#### Database Searching

The DIA raw files were processed using Spectronaut 18.0 in direct DIA mode^185^. We utilized a mouse (mus musculus, 25,435 protein entries) database comprising canonical and reviewed isoforms from Uniprot, supplemented with a list of 245 common protein contaminants and iRT peptides. Enzyme specificity was set to trypsin with allowance for two potential missed cleavages. Fixed modification was specified as carbamidomethyl of cysteine, while protein N-terminal acetylation and oxidation of methionine were considered variable modifications. To ensure high confidence, a false discovery rate limit of 1% was applied for precursors, peptides, and proteins identification, while the remaining search parameters were maintained at their default settings.

#### Bioinformatics Analysis

Proteomics data processing and statistical analysis were conducted in R. The MS2 intensity values generated by Spectronaut were utilized for analyzing the entire proteome dataset. The data underwent log2 transformation and normalization by subtracting the median value for each sample. To ensure data integrity, we filtered it to retain only proteins with complete values in at least one cohort. To compare proteomics data across groups, we employed a Limma t-test to identify proteins with differential abundance, and we visualized the impact of these differences through volcano plots. Lists of differentially abundant proteins were generated based on criteria of adjusted P.Value <0.05, resulting in a prioritized list for subsequent bioinformatics analysis.

### Statistical analysis

The statistical significance of our findings from flow cytometry, polysome fractionation, and Seahorse were assessed using unpaired t tests with Welch’s correction. We considered results with a p value less than 0.05 (p<0.05) to be statistically significant. * = p < 0.05, ** = p < 0.01, *** = p < 0.001. All error bars represent the standard error of the mean (SEM).

## Acknowledgments

We thank the Children’s Hospital Flow Cytometry Core, the Children’s Hospital of Philadelphia Proteomics Core Facility (RRID:SCR_023099), the Children’s Hospital of Philadelphia Center for Applied Genomics, and BC Children’s Hospital Research Institute Core Facilities for providing support and instrumentation; J. Henao-Mejia, P. Oliver, and C. Thaiss for feedback and thoughtful discussion; and all members of the Bailis laboratory for providing feedback and support.

## Author contributions

Conceptualization: MS, CSC, WB

Methodology: MS, MK, HF, LAS, CB, CM, RIKG, CSC, WB

Investigation: MS, KT, KR, ASA, JO, ET, ES, VL, CL, BG, CQ, LT, TP, JX

Formal analysis: MK

Visualization: MS, MK

Funding acquisition: WB

Project administration: WB

Supervision: CB, CM, RIKG, CSC, WB

Writing – original draft: MS, WB

Writing – review & editing: MS, RIKG, CSC, WB

## References Cited

1. Palm, W., and Thompson, C.B. (2017). Nutrient acquisition strategies of mammalian cells. Nature 546, 234–242. 10.1038/nature22379.

2. Zhu, J., and Thompson, C.B. (2019). Metabolic regulation of cell growth and proliferation. Nat Rev Mol Cell Biol 20, 436–450. 10.1038/s41580-019-0123-5.

3. Sung, Y., Yu, Y.C., and Han, J.M. (2023). Nutrient sensors and their crosstalk. Exp Mol Med 55, 1076–1089. 10.1038/s12276-023-01006-z.

4. Chantranupong, L., Wolfson, R.L., and Sabatini, D.M. (2015). Nutrient-sensing mechanisms across evolution. Cell 161, 67–83. 10.1016/j.cell.2015.02.041.

5. Efeyan, A., Comb, W.C., and Sabatini, D.M. (2015). Nutrient-sensing mechanisms and pathways. Nature 517, 302–310. 10.1038/nature14190.

6. Wek, R.C. (2018). Role of eIF2alpha Kinases in Translational Control and Adaptation to Cellular Stress. Cold Spring Harb Perspect Biol 10. 10.1101/cshperspect.a032870.

7. Wek, R.C., Anthony, T.G., and Staschke, K.A. (2023). Surviving and Adapting to Stress: Translational Control and the Integrated Stress Response. Antioxidants and Redox Signaling. Mary Ann Liebert Inc.

8. Pakos-Zebrucka, K., Koryga, I., Mnich, K., Ljujic, M., Samali, A., and Gorman, A.M. (2016). The integrated stress response. EMBO Rep. Springer Science and Business Media LLC.

9. Costa-Mattioli, M., and Walter, P. (2020). The integrated stress response: From mechanism to disease. Science (New York, N.Y.). NLM (Medline).

10. Gonzalez, A., and Hall, M.N. (2017). Nutrient sensing and TOR signaling in yeast and mammals. EMBO J 36, 397–408. 10.15252/embj.201696010.

11. Missiaen, R., Lesner, N.P., and Simon, M.C. (2023). HIF: a master regulator of nutrient availability and metabolic cross-talk in the tumor microenvironment. EMBO J 42, e112067. 10.15252/embj.2022112067.

12. Kamphorst, J.J., Nofal, M., Commisso, C., Hackett, S.R., Lu, W., Grabocka, E., Vander Heiden, M.G., Miller, G., Drebin, J.A., Bar-Sagi, D., et al. (2015). Human pancreatic cancer tumors are nutrient poor and tumor cells actively scavenge extracellular protein. Cancer Res 75, 544–553. 10.1158/0008-5472.CAN-14-2211.

13. Wellen, K.E., and Thompson, C.B. (2010). Cellular metabolic stress: considering how cells respond to nutrient excess. Mol Cell 40, 323–332. 10.1016/j.molcel.2010.10.004.

14. Elia, I., and Haigis, M.C. (2021). Metabolites and the tumour microenvironment: from cellular mechanisms to systemic metabolism. Nature Metabolism 2021 3:1. Nature Publishing Group.

15. Bantug, G.R., and Hess, C. (2023). The immunometabolic ecosystem in cancer. Nat Immunol 24, 2008–2020. 10.1038/s41590-023-01675-y.

16. Kao, K.C., Vilbois, S., Tsai, C.H., and Ho, P.C. (2022). Metabolic communication in the tumour-immune microenvironment. Nat Cell Biol 24, 1574–1583. 10.1038/s41556-022-01002-x.

17. Heintzman, D.R., Fisher, E.L., and Rathmell, J.C. (2022). Microenvironmental influences on T cell immunity in cancer and inflammation. Cellular & Molecular Immunology 2022 19:3. Nature Publishing Group.

18. Raynor, J.L., and Chi, H. (2024). Nutrients: Signal 4 in T cell immunity. J Exp Med 221. 10.1084/jem.20221839.

19. Franco, F., Jaccard, A., Romero, P., Yu, Y.R., and Ho, P.C. (2020). Metabolic and epigenetic regulation of T-cell exhaustion. Nat Metab 2, 1001–1012. 10.1038/s42255-020-00280-9.

20. Peng, J.J., Wang, L., Li, Z., Ku, C.L., and Ho, P.C. (2023). Metabolic challenges and interventions in CAR T cell therapy. Sci Immunol 8, eabq3016. 10.1126/sciimmunol.abq3016.

21. Wu, J., Li, G., Li, L., Li, D., Dong, Z., and Jiang, P. (2021). Asparagine enhances LCK signalling to potentiate CD8+ T-cell activation and anti-tumour responses. Nature Cell Biology 2021 23:1. Nature Publishing Group.

22. Gnanaprakasam, J.N.R., Kushwaha, B., Liu, L., Chen, X., Kang, S., Wang, T., Cassel, T.A., Adams, C.M., Higashi, R.M., Scott, D.A., et al. (2023). Asparagine restriction enhances CD8+ T cell metabolic fitness and antitumoral functionality through an NRF2-dependent stress response. Nature Metabolism 2023 5:8. Nature Publishing Group.

23. Vardhana, S.A., Hwee, M.A., Berisa, M., Wells, D.K., Yost, K.E., King, B., Smith, M., Herrera, P.S., Chang, H.Y., Satpathy, A.T., et al. (2020). Impaired mitochondrial oxidative phosphorylation limits the self-renewal of T cells exposed to persistent antigen. Nat Immunol 21, 1022–1033. 10.1038/s41590-020-0725-2.

24. Scharping, N.E., Rivadeneira, D.B., Menk, A.V., Vignali, P.D.A., Ford, B.R., Rittenhouse, N.L., Peralta, R., Wang, Y., Wang, Y., DePeaux, K., et al. (2021). Mitochondrial stress induced by continuous stimulation under hypoxia rapidly drives T cell exhaustion. Nature Immunology 2021 22:2. Nature Publishing Group.

25. Scharping, N.E., Menk, A.V., Moreci, R.S., Whetstone, R.D., Dadey, R.E., Watkins, S.C., Ferris, R.L., and Delgoffe, G.M. (2016). The Tumor Microenvironment Represses T Cell Mitochondrial Biogenesis to Drive Intratumoral T Cell Metabolic Insufficiency and Dysfunction. Immunity. Immunity.

26. Bell, H.N., Huber, A.K., Singhal, R., Korimerla, N., Rebernick, R.J., Kumar, R., El-derany, M.O., Sajjakulnukit, P., Das, N.K., Kerk, S.A., et al. (2023). Microenvironmental ammonia enhances T cell exhaustion in colorectal cancer. Cell Metabolism. Cell Press.

27. Bian, Y., Li, W., Kremer, D.M., Sajjakulnukit, P., Li, S., Crespo, J., Nwosu, Z.C., Zhang, L., Czerwonka, A., Pawłowska, A., et al. (2020). Cancer SLC43A2 alters T cell methionine metabolism and histone methylation. Nature. Nature.

28. Gemta, L.F., Siska, P.J., Nelson, M.E., Gao, X., Liu, X., Locasale, J.W., Yagita, H., Slingluff, C.L., Hoehn, K.L., Rathmell, J.C., and Bullock, T.N.J. (2019). Impaired enolase 1 glycolytic activity restrains effector functions of tumor-infiltrating CD8+ T cells. Science immunology. Sci Immunol.

29. Siska, P.J., Beckermann, K.E., Mason, F.M., Andrejeva, G., Greenplate, A.R., Sendor, A.B., Chiang, Y.C.J., Corona, A.L., Gemta, L.F., Vincent, B.G., et al. (2017). Mitochondrial dysregulation and glycolytic insufficiency functionally impair CD8 T cells infiltrating human renal cell carcinoma. JCI insight. JCI Insight.

30. Blagih, J., Coulombe, F., Vincent, E.E., Dupuy, F., Galicia-Vazquez, G., Yurchenko, E., Raissi, T.C., van der Windt, G.J., Viollet, B., Pearce, E.L., et al. (2015). The energy sensor AMPK regulates T cell metabolic adaptation and effector responses in vivo. Immunity 42, 41–54. 10.1016/j.immuni.2014.12.030.

31. Chang, C.H., Curtis, J.D., Maggi, L.B., Jr., Faubert, B., Villarino, A.V., O’Sullivan, D., Huang, S.C., van der Windt, G.J., Blagih, J., Qiu, J., et al. (2013). Posttranscriptional control of T cell effector function by aerobic glycolysis. Cell 153, 1239–1251. 10.1016/j.cell.2013.05.016.

32. Chang, C.H., Qiu, J., O’Sullivan, D., Buck, M.D., Noguchi, T., Curtis, J.D., Chen, Q., Gindin, M., Gubin, M.M., van der Windt, G.J., et al. (2015). Metabolic Competition in the Tumor Microenvironment Is a Driver of Cancer Progression. Cell 162, 1229–1241. 10.1016/j.cell.2015.08.016.

33. Qiu, J., Villa, M., Sanin, D.E., Buck, M.D., O’Sullivan, D., Ching, R., Matsushita, M., Grzes, K.M., Winkler, F., Chang, C.H., et al. (2019). Acetate Promotes T Cell Effector Function during Glucose Restriction. Cell Rep 27, 2063–2074 e2065. 10.1016/j.celrep.2019.04.022.

34. Yang, X., Xia, R., Yue, C., Zhai, W., Du, W., Yang, Q., Cao, H., Chen, X.X., Obando, D., Zhu, Y., et al. (2018). ATF4 Regulates CD4+ T Cell Immune Responses through Metabolic Reprogramming. Cell Reports. Cell Press.

35. Munn, D.H., Sharma, M.D., Baban, B., Harding, H.P., Zhang, Y., Ron, D., and Mellor, A.L. (2005). GCN2 kinase in T cells mediates proliferative arrest and anergy induction in response to indoleamine 2,3-dioxygenase. Immunity. Immunity.

36. Lu, Z., Bae, E.A., Verginadis, I.I., Zhang, H., Cho, C., McBrearty, N., George, S.S., Diehl, J.A., Koumenis, C., Bradley, L.M., and Fuchs, S.Y. (2023). Induction of the activating transcription factor-4 in the intratumoral CD8+ T cells sustains their viability and anti-tumor activities. Cancer immunology, immunotherapy : CII. Cancer Immunol Immunother.

37. Huang, H., Long, L., Zhou, P., Chapman, N.M., and Chi, H. (2020). mTOR signaling at the crossroads of environmental signals and T-cell fate decisions. Immunol Rev 295, 15–38. 10.1111/imr.12845.

38. Long, L., Wei, J., Lim, S.A., Raynor, J.L., Shi, H., Connelly, J.P., Wang, H., Guy, C., Xie, B., Chapman, N.M., et al. (2021). CRISPR screens unveil signal hubs for nutrient licensing of T cell immunity. Nature. Nature Research.

39. Huang, H., Zhou, P., Wei, J., Long, L., Shi, H., Dhungana, Y., Chapman, N.M., Fu, G., Saravia, J., Raynor, J.L., et al. (2021). In vivo CRISPR screening reveals nutrient signaling processes underpinning CD8+ T cell fate decisions. Cell.

40. Patel, C.H., Leone, R.D., Horton, M.R., and Powell, J.D. (2019). Targeting metabolism to regulate immune responses in autoimmunity and cancer. Nat Rev Drug Discov 18, 669–688. 10.1038/s41573-019-0032-5.

41. Wang, W., and Zou, W. (2020). Amino Acids and Their Transporters in T Cell Immunity and Cancer Therapy. Molecular Cell. Cell Press.

42. Yuan, H.X., Xiong, Y., and Guan, K.L. (2013). Nutrient sensing, metabolism, and cell growth control. Mol Cell 49, 379–387. 10.1016/j.molcel.2013.01.019.

43. Buccitelli, C., and Selbach, M. (2020). mRNAs, proteins and the emerging principles of gene expression control. Nat Rev Genet 21, 630–644. 10.1038/s41576-020-0258-4.

44. Philippe, L., van den Elzen, A.M.G., Watson, M.J., and Thoreen, C.C. (2020). Global analysis of LARP1 translation targets reveals tunable and dynamic features of 5’ TOP motifs. Proc Natl Acad Sci U S A 117, 5319–5328. 10.1073/pnas.1912864117.

45. Miloslavski, R., Cohen, E., Avraham, A., Iluz, Y., Hayouka, Z., Kasir, J., Mudhasani, R., Jones, S.N., Cybulski, N., Ruegg, M.A., et al. (2014). Oxygen sufficiency controls TOP mRNA translation via the TSC-Rheb-mTOR pathway in a 4E-BP-independent manner. J Mol Cell Biol 6, 255–266. 10.1093/jmcb/mju008.

46. Thoreen, C.C., Chantranupong, L., Keys, H.R., Wang, T., Gray, N.S., and Sabatini, D.M. (2012). A unifying model for mTORC1-mediated regulation of mRNA translation. Nature 485, 109–113. 10.1038/nature11083.

47. Roux, P.P., and Topisirovic, I. (2018). Signaling Pathways Involved in the Regulation of mRNA Translation. Mol Cell Biol 38. 10.1128/MCB.00070-18.

48. Araki, K., Morita, M., Bederman, A.G., Konieczny, B.T., Kissick, H.T., Sonenberg, N., and Ahmed, R. (2017). Translation is actively regulated during the differentiation of CD8(+) effector T cells. Nat Immunol. Nature Publishing Group.

49. Hsieh, A.C., Liu, Y., Edlind, M.P., Ingolia, N.T., Janes, M.R., Sher, A., Shi, E.Y., Stumpf, C.R., Christensen, C., Bonham, M.J., et al. (2012). The translational landscape of mTOR signalling steers cancer initiation and metastasis. Nature 485, 55–61. 10.1038/nature10912.

50. Koromilas, A.E. (2019). M(en)TORship lessons on life and death by the integrated stress response. Biochim Biophys Acta Gen Subj 1863, 644–649. 10.1016/j.bbagen.2018.12.009.

51. Han, S.H., Lee, M., Shin, Y., Giovanni, R., Chakrabarty, R.P., Herrerias, M.M., Dada, L.A., Flozak, A.S., Reyfman, P.A., Khuder, B., et al. (2023). Mitochondrial integrated stress response controls lung epithelial cell fate. Nature 2023 620:7975. Nature Publishing Group.

52. Liu, Q., Kang, S.A., Thoreen, C.C., Hur, W., Wang, J., Chang, J.W., Markhard, A., Zhang, J., Sim, T., Sabatini, D.M., and Gray, N.S. (2012). Development of ATP-competitive mTOR inhibitors. Methods Mol Biol 821, 447–460. 10.1007/978-1-61779-430-8_29.

53. Moerke, N.J., Aktas, H., Chen, H., Cantel, S., Reibarkh, M.Y., Fahmy, A., Gross, J.D., Degterev, A., Yuan, J., Chorev, M., et al. (2007). Small-molecule inhibition of the interaction between the translation initiation factors eIF4E and eIF4G. Cell 128, 257–267. 10.1016/j.cell.2006.11.046.

54. Keller, T.L., Zocco, D., Sundrud, M.S., Hendrick, M., Edenius, M., Yum, J., Kim, Y.J., Lee, H.K., Cortese, J.F., Wirth, D.F., et al. (2012). Halofuginone and other febrifugine derivatives inhibit prolyl-tRNA synthetase. Nat Chem Biol 8, 311–317. 10.1038/nchembio.790.

55. Ben-Sahra, I., and Manning, B.D. (2017). mTORC1 signaling and the metabolic control of cell growth. Curr. Opin. Cell Biol.

56. Torrence, M.E., Macarthur, M.R., Hosios, A.M., Valvezan, A.J., Asara, J.M., Mitchell, J.R., and Manning, B.D. (2021). The mtorc1-mediated activation of atf4 promotes protein and glutathione synthesis downstream of growth signals. eLife.

57. Valvezan, A.J., and Manning, B.D. (2019). Molecular logic of mTORC1 signalling as a metabolic rheostat. Nat. Metab.

58. Ben-Sahra, I., Hoxhaj, G., Ricoult, S.J.H., Asara, J.M., and Manning, B.D. (2016). mTORC1 induces purine synthesis through control of the mitochondrial tetrahydrofolate cycle. Science 351, 728–733. 10.1126/science.aad0489.

59. Brüggenthies, J.B., Fiore, A., Russier, M., Bitsina, C., Brötzmann, J., Kordes, S., Menninger, S., Wolf, A., Conti, E., Eickhoff, J.E., and Murray, P.J. (2022). A cell-based chemical-genetic screen for amino acid stress response inhibitors reveals torins reverse stress kinase GCN2 signaling. Journal of Biological Chemistry. American Society for Biochemistry and Molecular Biology Inc.

60. Cordova, R.A., Misra, J., Amin, P.H., Klunk, A.J., Damayanti, N.P., Carlson, K.R., Elmendorf, A.J., Kim, H.G., Mirek, E.T., Elzey, B.D., et al. (2022). GCN2 eIF2 kinase promotes prostate cancer by maintaining amino acid homeostasis. eLife. eLife Sciences Publications Ltd.

61. Nakamura, A., Nambu, T., Ebara, S., Hasegawa, Y., Toyoshima, K., Tsuchiya, Y., Tomita, D., Fujimoto, J., Kurasawa, O., Takahara, C., et al. (2018). Inhibition of GCN2 sensitizes ASNS-low cancer cells to asparaginase by disrupting the amino acid response. Proc Natl Acad Sci U S A 115, E7776–E7785. 10.1073/pnas.1805523115.

62. Huggins, C.J., Mayekar, M.K., Martin, N., Saylor, K.L., Gonit, M., Jailwala, P., Kasoji, M., Haines, D.C., Quiñones, O.A., and Johnson, P.F. C/EBPγ Is a Critical Regulator of Cellular Stress Response Networks through Heterodimerization with ATF4. Molecular and Cellular Biology.

63. Dey, S., Savant, S., Teske, B.F., Hatzoglou, M., Calkhoven, C.F., and Wek, R.C. (2012). Transcriptional repression of ATF4 gene by CCAAT/enhancer-binding protein beta (C/EBPbeta) differentially regulates integrated stress response. J Biol Chem 287, 21936–21949. 10.1074/jbc.M112.351783.

64. Kaspar, S., Oertlin, C., Szczepanowska, K., Kukat, A., Senft, K., Lucas, C., Brodesser, S., Hatzoglou, M., Larsson, O., Topisirovic, I., and Trifunovic, A. (2021). Adaptation to mitochondrial stress requires CHOP-directed tuning of ISR. Sci Adv 7. 10.1126/sciadv.abf0971.

65. Ma, E.H., Bantug, G., Griss, T., Condotta, S., Johnson, R.M., Samborska, B., Mainolfi, N., Suri, V., Guak, H., Balmer, M.L., et al. (2017). Serine Is an Essential Metabolite for Effector T Cell Expansion. Cell Metabolism. Cell Press.

66. Ma, E.H., Verway, M.J., Johnson, R.M., Roy, D.G., Steadman, M., Hayes, S., Williams, K.S., Sheldon, R.D., Samborska, B., Kosinski, P.A., et al. (2019). Metabolic Profiling Using Stable Isotope Tracing Reveals Distinct Patterns of Glucose Utilization by Physiologically Activated CD8(+) T Cells. Immunity 51, 856–870 e855. 10.1016/j.immuni.2019.09.003.

67. Ron-Harel, N., Santos, D., Ghergurovich, J.M., Sage, P.T., Reddy, A., Lovitch, S.B., Dephoure, N., Satterstrom, F.K., Sheffer, M., Spinelli, J.B., et al. (2016). Mitochondrial biogenesis and proteome remodeling promote one-carbon metabolism for T cell activation. Cell Metab. Cell Press.

68. Ron-Harel, N., Ghergurovich, J.M., Notarangelo, G., LaFleur, M.W., Tsubosaka, Y., Sharpe, A.H., Rabinowitz, J.D., and Haigis, M.C. (2019). T Cell Activation Depends on Extracellular Alanine. Cell reports. Cell Rep.

69. Monod, J. (1949). THE GROWTH OF BACTERIAL CULTURES. Annual Review of Microbiology 3, 371–394. 10.1146/annurev.mi.03.100149.002103.

70. Jacob, F., and Monod, J. (1961). Genetic regulatory mechanisms in the synthesis of proteins. J Mol Biol 3, 318–356. 10.1016/s0022-2836(61)80072-7.

71. McKnight, S.L. (2010). On getting there from here. Science 330, 1338–1339. 10.1126/science.1199908.

72. Blaiseau, P.L., and Holmes, A.M. (2021). Diauxic Inhibition: Jacques Monod’s Ignored Work. J Hist Biol 54, 175–196. 10.1007/s10739-021-09639-4.

73. Bagamery, L.E., Justman, Q.A., Garner, E.C., and Murray, A.W. (2020). A Putative Bet-Hedging Strategy Buffers Budding Yeast against Environmental Instability. Curr Biol 30, 4563–4578 e4564. 10.1016/j.cub.2020.08.092.

74. Conrad, M., Schothorst, J., Kankipati, H.N., Van Zeebroeck, G., Rubio-Texeira, M., and Thevelein, J.M. (2014). Nutrient sensing and signaling in the yeast Saccharomyces cerevisiae. FEMS Microbiol Rev 38, 254–299. 10.1111/1574-6976.12065.

75. Wang, J., Atolia, E., Hua, B., Savir, Y., Escalante-Chong, R., and Springer, M. (2015). Natural variation in preparation for nutrient depletion reveals a cost-benefit tradeoff. PLoS Biol 13, e1002041. 10.1371/journal.pbio.1002041.

76. Gorke, B., and Stulke, J. (2008). Carbon catabolite repression in bacteria: many ways to make the most out of nutrients. Nat Rev Microbiol 6, 613–624. 10.1038/nrmicro1932.

77. Pavlova, N.N., and Thompson, C.B. (2024). Oncogenic Control of Metabolism. Cold Spring Harb Perspect Med 14. 10.1101/cshperspect.a041531.

78. Pavlova, N.N., Zhu, J., and Thompson, C.B. (2022). The hallmarks of cancer metabolism: Still emerging. Cell Metab 34, 355–377. 10.1016/j.cmet.2022.01.007.

79. Ma, E.H., Dahabieh, M.S., DeCamp, L.M., Kaymak, I., Kitchen-Goosen, S.M., Oswald, B.M., Longo, J., Roy, D.G., Verway, M.J., Johnson, R.M., et al. (2024). (13)C metabolite tracing reveals glutamine and acetate as critical in vivo fuels for CD8 T cells. Sci Adv 10, eadj1431. 10.1126/sciadv.adj1431.

80. Wang, T., Gnanaprakasam, J.N.R., Chen, X., Kang, S., Xu, X., Sun, H., Liu, L., Rodgers, H., Miller, E., Cassel, T.A., et al. (2020). Inosine is an alternative carbon source for CD8(+)-T-cell function under glucose restriction. Nat Metab 2, 635–647. 10.1038/s42255-020-0219-4.

81. Kaymak, I., Luda, K.M., Duimstra, L.R., Ma, E.H., Longo, J., Dahabieh, M.S., Faubert, B., Oswald, B.M., Watson, M.J., Kitchen-Goosen, S.M., et al. (2022). Carbon source availability drives nutrient utilization in CD8(+) T cells. Cell Metab 34, 1298–1311 e1296. 10.1016/j.cmet.2022.07.012.

82. Luda, K.M., Longo, J., Kitchen-Goosen, S.M., Duimstra, L.R., Ma, E.H., Watson, M.J., Oswald, B.M., Fu, Z., Madaj, Z., Kupai, A., et al. (2023). Ketolysis drives CD8(+) T cell effector function through effects on histone acetylation. Immunity 56, 2021–2035 e2028. 10.1016/j.immuni.2023.07.002.

83. van der Windt, G.J., and Pearce, E.L. (2012). Metabolic switching and fuel choice during T-cell differentiation and memory development. Immunol Rev 249, 27–42. 10.1111/j.1600-065X.2012.01150.x.

84. Fox, C.J., Hammerman, P.S., and Thompson, C.B. (2005). Fuel feeds function: energy metabolism and the T-cell response. Nat Rev Immunol 5, 844–852. 10.1038/nri1710.

85. Hui, S., Ghergurovich, J.M., Morscher, R.J., Jang, C., Teng, X., Lu, W., Esparza, L.A., Reya, T., Le, Z., Yanxiang Guo, J., et al. (2017). Glucose feeds the TCA cycle via circulating lactate. Nature 551, 115–118. 10.1038/nature24057.

86. Zhang, Y., Kurupati, R., Liu, L., Zhou, X.Y., Zhang, G., Hudaihed, A., Filisio, F., Giles-Davis, W., Xu, X., Karakousis, G.C., et al. (2017). Enhancing CD8+ T Cell Fatty Acid Catabolism within a Metabolically Challenging Tumor Microenvironment Increases the Efficacy of Melanoma Immunotherapy. Cancer cell. Cancer Cell.

87. Shen, Y., Dinh, H.V., Cruz, E.R., Chen, Z., Bartman, C.R., Xiao, T., Call, C.M., Ryseck, R.P., Pratas, J., Weilandt, D., et al. (2024). Mitochondrial ATP generation is more proteome efficient than glycolysis. Nat Chem Biol 20, 1123–1132. 10.1038/s41589-024-01571-y.

88. Chen, C., Zheng, H., Horwitz, E.M., Ando, S., Araki, K., Zhao, P., Li, Z., Ford, M.L., Ahmed, R., and Qu, C.K. (2023). Mitochondrial metabolic flexibility is critical for CD8(+) T cell antitumor immunity. Sci Adv 9, eadf9522. 10.1126/sciadv.adf9522.

89. Condon, K.J., Orozco, J.M., Adelmann, C.H., Spinelli, J.B., van der Helm, P.W., Roberts, J.M., Kunchok, T., and Sabatini, D.M. (2021). Genome-wide CRISPR screens reveal multitiered mechanisms through which mTORC1 senses mitochondrial dysfunction. Proceedings of the National Academy of Sciences of the United States of America. National Academy of Sciences.

90. Park, Y., Reyna-Neyra, A., Philippe, L., and Thoreen, C.C. (2017). mTORC1 Balances Cellular Amino Acid Supply with Demand for Protein Synthesis through Post-transcriptional Control of ATF4. Cell Reports. Cell Press.

91. Misra, J., Holmes, M.J., Mirek, E.T., Langevin, M., Kim, H.G., Carlson, K.R., Watford, M., Charlie Dong, X., Anthony, T.G., and Wek, R.C. (2021). Discordant regulation of eIF2 kinase GCN2 and mTORC1 during nutrient stress.

92. Ye, J., Palm, W., Peng, M., King, B., Lindsten, T., Li, M.O., and al., e. (2015). GCN2 sustains mTORC1 suppression upon amino acid deprivation by inducing Sestrin2. Genes Dev.

93. Darawshi, O., Yassin, O., Shmuel, M., Wek, R.C., Mahdizadeh, S.J., Eriksson, L.A., Hatzoglou, M., and Tirosh, B. (2024). Phosphorylation of GCN2 by mTOR confers adaptation to conditions of hyper-mTOR activation under stress. J Biol Chem 300, 107575. 10.1016/j.jbc.2024.107575.

94. Neill, G., and Masson, G.R. (2023). A stay of execution: ATF4 regulation and potential outcomes for the integrated stress response. Frontiers in Molecular Neuroscience. Frontiers Media S.A.

95. Pollizzi, K.N., and Powell, J.D. (2015). Regulation of T cells by mTOR: the known knowns and the known unknowns. Trends Immunol 36, 13–20. 10.1016/j.it.2014.11.005.

96. Carlson, T.J., Pellerin, A., Djuretic, I.M., Trivigno, C., Koralov, S.B., Rao, A., and Sundrud, M.S. (2014). Halofuginone-Induced Amino Acid Starvation Regulates Stat3-Dependent Th17 Effector Function and Reduces Established Autoimmune Inflammation. Journal of Immunology. The American Association of Immunologists.

97. Cao, Y., Trillo-Tinoco, J., Sierra, R.A., Anadon, C., Dai, W., Mohamed, E., Cen, L., Costich, T.L., Magliocco, A., Marchion, D., et al. (2019). ER stress-induced mediator C/EBP homologous protein thwarts effector T cell activity in tumors through T-bet repression. Nature Communications 2019 10:1. Nature Publishing Group.

98. Zou, Z., Cheng, Q., Zhou, J., Guo, C., Hadjinicolaou, A.V., Salio, M., Liang, X., Yang, C., Du, Y., Yao, W., et al. (2024). ATF4-SLC7A11-GSH axis mediates the acquisition of immunosuppressive properties by activated CD4+ T cells in low arginine condition. Cell Reports. Elsevier B.V.

99. Di Conza, G., Ho, P.C., Cubillos-Ruiz, J.R., and Huang, S.C.C. (2023). Control of immune cell function by the unfolded protein response.

100. Rashidi, A., Miska, J., Lee-Chang, C., Kanojia, D., Panek, W.K., Lopez-Rosas, A., Zhang, P., Han, Y., Xiao, T., Pituch, K.C., et al. (2020). GCN2 is essential for CD8+ T cell survival and function in murine models of malignant glioma. Cancer Immunology, Immunotherapy. Springer.

101. Scheu, S., Stetson, D.B., Reinhardt, R.L., Leber, J.H., Mohrs, M., and Locksley, R.M. (2006). Activation of the integrated stress response during T helper cell differentiation. Nature Immunology 2006 7:6. Nature Publishing Group.

102. St. Paul, M., Saibil, S.D., Kates, M., Han, S., Lien, S.C., Laister, R.C., Hezaveh, K., Kloetgen, A., Penny, S., Guo, T., et al. (2024). Ex vivo activation of the GCN2 pathway metabolically reprograms T cells, leading to enhanced adoptive cell therapy. Cell Reports Medicine. Cell Press.

103. Jurgens, A.P., Popović, B., and Wolkers, M.C. (2021). T cells at work: How post-transcriptional mechanisms control T cell homeostasis and activation. European Journal of Immunology. John Wiley & Sons, Ltd.

104. Salerno, F., Engels, S., van den Biggelaar, M., van Alphen, F.P.J., Guislain, A., Zhao, W., Hodge, D.L., Bell, S.E., Medema, J.P., von Lindern, M., et al. (2018). Translational repression of pre-formed cytokine-encoding mRNA prevents chronic activation of memory T cells. Nature Immunology 2018 19:8. Nature Publishing Group.

105. Riesenberg, B.P., Hunt, E.G., Tennant, M.D., Hurst, K.E., Andrews, A.M., Leddy, L.R., Neskey, D.M., Hill, E.G., Rivera, G.O.R., Paulos, C.M., et al. (2022). Stress-Mediated Attenuation of Translation Undermines T-cell Activity in Cancer. Cancer Research. American Association for Cancer Research Inc.

106. Wolf, T., Jin, W., Zoppi, G., Vogel, I.A., Akhmedov, M., Bleck, C.K.E., Beltraminelli, T., Rieckmann, J.C., Ramirez, N.J., Benevento, M., et al. (2020). Dynamics in protein translation sustaining T cell preparedness. Nat Immunol. Nature Research.

107. Tan, T.C.J., Knight, J., Sbarrato, T., Dudek, K., Willis, A.E., and Zamoyska, R. (2017). Suboptimal T-cell receptor signaling compromises protein translation, ribosome biogenesis, and proliferation of mouse CD8 T cells. Proc Natl Acad Sci USA. National Academy of Sciences.

108. Tan, T.C.J., Kelly, V., Zou, X., Wright, D., Ly, T., and Zamoyska, R. (2022). Translation factor eIF5a is essential for IFNγ production and cell cycle regulation in primary CD8+ T lymphocytes. Nature Communications 2022 13:1. Nature Publishing Group.

109. Ricciardi, S., Manfrini, N., Alfieri, R., Calamita, P., Crosti, M.C., Gallo, S., Müller, R., Pagani, M., Abrignani, S., and Biffo, S. (2018). The Translational Machinery of Human CD4(+) T Cells Is Poised for Activation and Controls the Switch from Quiescence to Metabolic Remodeling. Cell Metab. Cell Press.

110. Lisci, M., Barton, P.R., Randzavola, L.O., Ma, C.Y., Marchingo, J.M., Cantrell, D.A., Paupe, V., Prudent, J., Stinchcombe, J.C., and Griffiths, G.M. (2021). Mitochondrial translation is required for sustained killing by cytotoxic T cells. Science. American Association for the Advancement of Science.

111. Liedmann, S., Liu, X., Guy, C.S., Crawford, J.C., Rodriguez, D.A., Kuzuoglu-Ozturk, D., Guo, A., Verbist, K.C., Temirov, J., Chen, M.J., et al. (2022). Localization of a TORC1-eIF4F translation complex during CD8(+) T cell activation drives divergent cell fate. Mol Cell 82, 2401–2414 e2409. 10.1016/j.molcel.2022.04.016.

112. Bjur, E., Larsson, O., Yurchenko, E., Zheng, L., Gandin, V., Topisirovic, I., Li, S., Wagner, C.R., Sonenberg, N., and Piccirillo, C.A. (2013). Distinct translational control in CD4+ T cell subsets. PLoS Genet.

113. Piccirillo, C.A., Bjur, E., Topisirovic, I., Sonenberg, N., and Larsson, O. (2014). Translational control of immune responses: from transcripts to translatomes. Nature Publishing Group.

114. Giles, J.R., Ngiow, S.F., Manne, S., Baxter, A.E., Khan, O., Wang, P., Staupe, R., Abdel-Hakeem, M.S., Huang, H., Mathew, D., et al. (2022). Shared and distinct biological circuits in effector, memory and exhausted CD8(+) T cells revealed by temporal single-cell transcriptomics and epigenetics. Nat Immunol 23, 1600–1613. 10.1038/s41590-022-01338-4.

115. Hwang, S.S., Lim, J., Yu, Z., Kong, P., Sefik, E., Xu, H., Harman, C.C.D., Kim, L.K., Lee, G.R., Li, H.B., and Flavell, R.A. (2020). mRNA destabilization by BTG1 and BTG2 maintains T cell quiescence. Science 367, 1255–1260. 10.1126/science.aax0194.

116. Yuniati, L., Scheijen, B., van der Meer, L.T., and van Leeuwen, F.N. (2019). Tumor suppressors BTG1 and BTG2: Beyond growth control. J Cell Physiol 234, 5379–5389. 10.1002/jcp.27407.

117. Liu, X., Wang, Y., Lu, H., Li, J., Yan, X., Xiao, M., Hao, J., Alekseev, A., Khong, H., Chen, T., et al. (2019). Genome-wide analysis identifies NR4A1 as a key mediator of T cell dysfunction. Nature 567, 525–529. 10.1038/s41586-019-0979-8.

118. Chen, J., Lopez-Moyado, I.F., Seo, H., Lio, C.J., Hempleman, L.J., Sekiya, T., Yoshimura, A., Scott-Browne, J.P., and Rao, A. (2019). NR4A transcription factors limit CAR T cell function in solid tumours. Nature 567, 530–534. 10.1038/s41586-019-0985-x.

119. Peralta, R.M., Xie, B., Lontos, K., Nieves-Rosado, H., Spahr, K., Joshi, S., Ford, B.R., Quann, K., Frisch, A.T., Dean, V., et al. (2024). Dysfunction of exhausted T cells is enforced by MCT11-mediated lactate metabolism. Nat Immunol 25, 2297–2307. 10.1038/s41590-024-01999-3.

120. Bengsch, B., Johnson, A.L., Kurachi, M., Odorizzi, P.M., Pauken, K.E., Attanasio, J., Stelekati, E., McLane, L.M., Paley, M.A., Delgoffe, G.M., and Wherry, E.J. (2016). Bioenergetic Insufficiencies Due to Metabolic Alterations Regulated by the Inhibitory Receptor PD-1 Are an Early Driver of CD8(+) T Cell Exhaustion. Immunity 45, 358–373. 10.1016/j.immuni.2016.07.008.

121. Wu, H., Zhao, X., Hochrein, S.M., Eckstein, M., Gubert, G.F., Knöpper, K., Mansilla, A.M., Öner, A., Doucet-Ladevèze, R., Schmitz, W., et al. (2023). Mitochondrial dysfunction promotes the transition of precursor to terminally exhausted T cells through HIF-1α-mediated glycolytic reprogramming. Nature Communications 2023 14:1. Nature Publishing Group.

122. Zebley, C.C., Zehn, D., Gottschalk, S., and Chi, H. (2024). T cell dysfunction and therapeutic intervention in cancer. Nat Immunol 25, 1344–1354. 10.1038/s41590-024-01896-9.

123. Salerno, F., Paolini, N.A., Stark, R., Von Lindern, M., and Wolkers, M.C. (2017). Distinct PKC-mediated posttranscriptional events set cytokine production kinetics in CD8+ T cells. Proceedings of the National Academy of Sciences of the United States of America. National Academy of Sciences.

124. Salerno, F., Freen-van Heeren, J.J., Guislain, A., Nicolet, B.P., and Wolkers, M.C. (2019). Costimulation through TLR2 Drives Polyfunctional CD8+ T Cell Responses. The Journal of Immunology. American Association of Immunologists.

125. Salerno, F., Turner, M., and Wolkers, M.C. (2020). Dynamic Post-Transcriptional Events Governing CD8+ T Cell Homeostasis and Effector Function. Trends in Immunology. Elsevier Current Trends.

126. Popovic, B., Nicolet, B.P., Guislain, A., Engels, S., Jurgens, A.P., Paravinja, N., Freen-van Heeren, J.J., van Alphen, F.P.J., van den Biggelaar, M., Salerno, F., and Wolkers, M.C. (2023). Time-dependent regulation of cytokine production by RNA binding proteins defines T cell effector function. Cell Rep 42, 112419. 10.1016/j.celrep.2023.112419.

127. Freen-van Heeren, J.J., Popović, B., Guislain, A., and Wolkers, M.C. (2020). Human T cells employ conserved AU-rich elements to fine-tune IFN-γ production. European Journal of Immunology. John Wiley & Sons, Ltd.

128. Nicolet, B.P., and Wolkers, M.C. (2022). The relationship of mRNA with protein expression in CD8+ T cells associates with gene class and gene characteristics. PLOS ONE. Public Library of Science.

129. Leibovitch, M., and Topisirovic, I. (2018). Dysregulation of mRNA translation and energy metabolism in cancer. Adv Biol Regul 67, 30–39. 10.1016/j.jbior.2017.11.001.

130. Lindqvist, L.M., Tandoc, K., Topisirovic, I., and Furic, L. (2018). Cross-talk between protein synthesis, energy metabolism and autophagy in cancer. Curr Opin Genet Dev 48, 104–111. 10.1016/j.gde.2017.11.003.

131. Orr, M.W., Mao, Y., Storz, G., and Qian, S.B. (2020). Alternative ORFs and small ORFs: shedding light on the dark proteome. Nucleic Acids Res 48, 1029–1042. 10.1093/nar/gkz734.

132. Leppek, K., Das, R., and Barna, M. (2018). Functional 5’ UTR mRNA structures in eukaryotic translation regulation and how to find them. Nat Rev Mol Cell Biol 19, 158–174. 10.1038/nrm.2017.103.

133. Adamson, B., Norman, T.M., Jost, M., Cho, M.Y., Nuñez, J.K., Chen, Y., Villalta, J.E., Gilbert, L.A., Horlbeck, M.A., Hein, M.Y., et al. (2016). A Multiplexed Single-Cell CRISPR Screening Platform Enables Systematic Dissection of the Unfolded Protein Response. Cell. Cell Press.

134. Replogle, J.M., Norman, T.M., Xu, A., Hussmann, J.A., Chen, J., Cogan, J.Z., Meer, E.J., Terry, J.M., Riordan, D.P., Srinivas, N., et al. (2020). Combinatorial single-cell CRISPR screens by direct guide RNA capture and targeted sequencing. Nature Biotechnology 2020 38:8. Nature Publishing Group.

135. Shi, H., Doench, J.G., and Chi, H. (2023). CRISPR screens for functional interrogation of immunity. Nat Rev Immunol 23, 363–380. 10.1038/s41577-022-00802-4.

136. Halaby, M.J., Hezaveh, K., Lamorte, S., Ciudad, M.T., Kloetgen, A., MacLeod, B.L., Guo, M., Chakravarthy, A., Da Silva Medina, T., Ugel, S., et al. (2019). GCN2 drives macrophage and MDSC function and immunosuppression in the tumor microenvironment. Science Immunology. American Association for the Advancement of Science.

137. Ravindran, R., Khan, N., Nakaya, H.I., Li, S., Loebbermann, J., Maddur, M.S., Park, Y., Jones, D.P., Chappert, P., Davoust, J., et al. (2014). Vaccine activation of the nutrient sensor GCN2 in dendritic cells enhances antigen presentation. Science (New York, N.Y.). Science.

138. Yazicioglu, Y.F., Marin, E., Andrew, H.F., Bentkowska, K., Johnstone, J.C., Mitchell, R., Wong, Z.Y., Zec, K., Fergusson, J., Borsa, M., et al. (2024). Asparagine availability controls germinal center B cell homeostasis. Sci Immunol 9, eadl4613. 10.1126/sciimmunol.adl4613.

139. Ravindran, R., Loebbermann, J., Nakaya, H.I., Khan, N., Ma, H., Gama, L., Machiah, D.K., Lawson, B., Hakimpour, P., Wang, Y.C., et al. (2016). The amino acid sensor GCN2 controls gut inflammation by inhibiting inflammasome activation. Nature 2016 531:7595. Nature Publishing Group.

140. Fletcher, M., Ramirez, M.E., Sierra, R.A., Raber, P., Thevenot, P., Al-Khami, A.A., Sanchez-Pino, D., Hernez, C., Wyczechowska, D.D., Ochoa, A.C., and Rodriguez, P.C. (2015). l-Arginine depletion blunts antitumor T-cell responses by inducing myeloid-derived suppressor cells. Cancer research. Cancer Res.

141. Van De Velde, L.A., Subramanian, C., Smith, A.M., Barron, L., Qualls, J.E., Neale, G., Alfonso-Pecchio, A., Jackowski, S., Rock, C.O., Wynn, T.A., and Murray, P.J. (2017). T Cells Encountering Myeloid Cells Programmed for Amino Acid-dependent Immunosuppression Use Rictor/mTORC2 Protein for Proliferative Checkpoint Decisions. Journal of Biological Chemistry. Elsevier.

142. Klein Geltink, R.I., Edwards-Hicks, J., Apostolova, P., O’Sullivan, D., Sanin, D.E., Patterson, A.E., Puleston, D.J., Ligthart, N.A.M., Buescher, J.M., Grzes, K.M., et al. (2020). Metabolic conditioning of CD8(+) effector T cells for adoptive cell therapy. Nat Metab 2, 703–716. 10.1038/s42255-020-0256-z.

143. Geiger, R., Rieckmann, J.C., Wolf, T., Basso, C., Feng, Y., Fuhrer, T., Kogadeeva, M., Picotti, P., Meissner, F., Mann, M., et al. (2016). L-Arginine Modulates T Cell Metabolism and Enhances Survival and Anti-tumor Activity. Cell.

144. Kishton, R.J., Sukumar, M., and Restifo, N.P. (2017). Metabolic Regulation of T Cell Longevity and Function in Tumor Immunotherapy. Cell Metab 26, 94–109. 10.1016/j.cmet.2017.06.016.

145. Mishra, D., and Banerjee, D. (2021). Metabolic Interactions Between Tumor and Stromal Cells in the Tumor Microenvironment. Adv Exp Med Biol 1350, 101–121. 10.1007/978-3-030-83282-7_5.

146. Lyssiotis, C.A., and Kimmelman, A.C. (2017). Metabolic Interactions in the Tumor Microenvironment. Trends Cell Biol 27, 863–875. 10.1016/j.tcb.2017.06.003.

147. Sela, Y., Li, J., Maheswaran, S., Norgard, R., Yuan, S., Hubbi, M., Doepner, M., Xu, J.P., Ho, E.S., Mesaros, C., et al. (2022). Bcl-xL Enforces a Slow-Cycling State Necessary for Survival in the Nutrient-Deprived Microenvironment of Pancreatic Cancer. Cancer Res 82, 1890–1908. 10.1158/0008-5472.CAN-22-0431.

148. Vander Heiden, M.G., and DeBerardinis, R.J. (2017). Understanding the intersections between metabolism and cancer biology. Cell. NIH Public Access.

149. Muir, A., Danai, L.V., Gui, D.Y., Waingarten, C.Y., Lewis, C.A., and Vander Heiden, M.G. (2017). Environmental cystine drives glutamine anaplerosis and sensitizes cancer cells to glutaminase inhibition. eLife. eLife Sciences Publications Ltd.

150. Gui, D.Y., Sullivan, L.B., Luengo, A., Hosios, A.M., Bush, L.N., Gitego, N., Davidson, S.M., Freinkman, E., Thomas, C.J., and Vander Heiden, M.G. (2016). Environment Dictates Dependence on Mitochondrial Complex I for NAD+ and Aspartate Production and Determines Cancer Cell Sensitivity to Metformin. Cell metabolism. Cell Metab.

151. Davidson, S.M., Papagiannakopoulos, T., Olenchock, B.A., Heyman, J.E., Keibler, M.A., Luengo, A., Bauer, M.R., Jha, A.K., O’Brien, J.P., Pierce, K.A., et al. (2016). Environment impacts the metabolic dependencies of Ras-driven non-small cell lung cancer. Cell Metab. Cell Press.

152. Fabbri, L., Chakraborty, A., Robert, C., and Vagner, S. (2021). The plasticity of mRNA translation during cancer progression and therapy resistance. Nature Reviews Cancer 2021 21:9. Nature Publishing Group.

153. Franca, G.S., Baron, M., King, B.R., Bossowski, J.P., Bjornberg, A., Pour, M., Rao, A., Patel, A.S., Misirlioglu, S., Barkley, D., et al. (2024). Cellular adaptation to cancer therapy along a resistance continuum. Nature 631, 876–883. 10.1038/s41586-024-07690-9.

154. Hulea, L., Gravel, S.P., Morita, M., Cargnello, M., Uchenunu, O., Im, Y.K., Lehuede, C., Ma, E.H., Leibovitch, M., McLaughlan, S., et al. (2018). Translational and HIF-1alpha-Dependent Metabolic Reprogramming Underpin Metabolic Plasticity and Responses to Kinase Inhibitors and Biguanides. Cell Metab 28, 817–832 e818. 10.1016/j.cmet.2018.09.001.

155. Gaude, E., and Frezza, C. (2016). Tissue-specific and convergent metabolic transformation of cancer correlates with metastatic potential and patient survival. Nature Communications 2016 7:1. Nature Publishing Group.

156. Ghaddar, N., Wang, S., Woodvine, B., Krishnamoorthy, J., van Hoef, V., Darini, C., Kazimierczak, U., Ah-Son, N., Popper, H., Johnson, M., et al. (2021). The integrated stress response is tumorigenic and constitutes a therapeutic liability in KRAS-driven lung cancer. Nat Commun 12, 4651. 10.1038/s41467-021-24661-0.

157. Gwinn, D.M., Lee, A.G., Briones-Martin-del-Campo, M., Conn, C.S., Simpson, D.R., Scott, A.I., Le, A., Cowan, T.M., Ruggero, D., and Sweet-Cordero, E.A. (2018). Oncogenic KRAS Regulates Amino Acid Homeostasis and Asparagine Biosynthesis via ATF4 and Alters Sensitivity to L-Asparaginase. Cancer Cell. Cell Press.

158. Vasan, N., Baselga, J., and Hyman, D.M. (2019). A view on drug resistance in cancer. Nature 575, 299–309. 10.1038/s41586-019-1730-1.

159. Halbrook, C.J., Thurston, G., Boyer, S., Anaraki, C., Jimenez, J.A., McCarthy, A., Steele, N.G., Kerk, S.A., Hong, H.S., Lin, L., et al. (2022). Differential integrated stress response and asparagine production drive symbiosis and therapy resistance of pancreatic adenocarcinoma cells. Nat Cancer 3, 1386–1403. 10.1038/s43018-022-00463-1.

160. Kerk, S.A., Papagiannakopoulos, T., Shah, Y.M., and Lyssiotis, C.A. (2021). Metabolic networks in mutant KRAS-driven tumours: tissue specificities and the microenvironment. Nature Reviews Cancer 2021 21:8. Nature Publishing Group.

161. Marine, J.C., Dawson, S.J., and Dawson, M.A. (2020). Non-genetic mechanisms of therapeutic resistance in cancer. Nat Rev Cancer 20, 743–756. 10.1038/s41568-020-00302-4.

162. Ye, J., Kumanova, M., Hart, L.S., Sloane, K., Zhang, H., De Panis, D.N., Bobrovnikova-Marjon, E., Diehl, J.A., Ron, D., and Koumenis, C. (2010). The GCN2-ATF4 pathway is critical for tumour cell survival and proliferation in response to nutrient deprivation. EMBO J 29, 2082–2096. 10.1038/emboj.2010.81.

163. Andrews, S. (2010). FASTQC. A quality control tool for high throughput sequence data.

164. Ewels, P., Magnusson, M., Lundin, S., and Kaller, M. (2016). MultiQC: summarize analysis results for multiple tools and samples in a single report. Bioinformatics 32, 3047–3048. 10.1093/bioinformatics/btw354.

165. Bolger, A.M., Lohse, M., and Usadel, B. (2014). Trimmomatic: a flexible trimmer for Illumina sequence data. Bioinformatics 30, 2114–2120. 10.1093/bioinformatics/btu170.

166. Mouse Genome Sequencing, C., Waterston, R.H., Lindblad-Toh, K., Birney, E., Rogers, J., Abril, J.F., Agarwal, P., Agarwala, R., Ainscough, R., Alexandersson, M., et al. (2002). Initial sequencing and comparative analysis of the mouse genome. Nature 420, 520–562. 10.1038/nature01262.

167. Frankish, A., Diekhans, M., Jungreis, I., Lagarde, J., Loveland, J.E., Mudge, J.M., Sisu, C., Wright, J.C., Armstrong, J., Barnes, I., et al. (2021). Gencode 2021. Nucleic Acids Res 49, D916–D923. 10.1093/nar/gkaa1087.

168. Dobin, A., Davis, C.A., Schlesinger, F., Drenkow, J., Zaleski, C., Jha, S., Batut, P., Chaisson, M., and Gingeras, T.R. (2013). STAR: ultrafast universal RNA-seq aligner. Bioinformatics 29, 15–21. 10.1093/bioinformatics/bts635.

169. Li, H., Handsaker, B., Wysoker, A., Fennell, T., Ruan, J., Homer, N., Marth, G., Abecasis, G., Durbin, R., and Genome Project Data Processing, S. (2009). The Sequence Alignment/Map format and SAMtools. Bioinformatics 25, 2078–2079. 10.1093/bioinformatics/btp352.

170. Anders, S., Pyl, P.T., and Huber, W. (2015). HTSeq--a Python framework to work with high-throughput sequencing data. Bioinformatics 31, 166–169. 10.1093/bioinformatics/btu638.

171. Robinson, M.D., McCarthy, D.J., and Smyth, G.K. (2010). edgeR: a Bioconductor package for differential expression analysis of digital gene expression data. Bioinformatics 26, 139–140. 10.1093/bioinformatics/btp616.

172. Durinck, S., Spellman, P.T., Birney, E., and Huber, W. (2009). Mapping identifiers for the integration of genomic datasets with the R/Bioconductor package biomaRt. Nat Protoc 4, 1184–1191. 10.1038/nprot.2009.97.

173. Oertlin, C., Lorent, J., Murie, C., Furic, L., Topisirovic, I., and Larsson, O. (2019). Generally applicable transcriptome-wide analysis of translation using anota2seq. Nucleic Acids Res 47, e70. 10.1093/nar/gkz223.

174. Wickham, H. (2011). ggplot2. WIREs Computational Statistics 3, 180–185. 10.1002/wics.147.

175. Conway, J.R., Lex, A., and Gehlenborg, N. (2017). UpSetR: an R package for the visualization of intersecting sets and their properties. Bioinformatics 33, 2938–2940. 10.1093/bioinformatics/btx364.

176. Kolberg, L., Raudvere, U., Kuzmin, I., Vilo, J., and Peterson, H. (2020). gprofiler2 -- an R package for gene list functional enrichment analysis and namespace conversion toolset g:Profiler. F1000Res 9. 10.12688/f1000research.24956.2.

177. Gu, Z., Eils, R., and Schlesner, M. (2016). Complex heatmaps reveal patterns and correlations in multidimensional genomic data. Bioinformatics 32, 2847–2849. 10.1093/bioinformatics/btw313.

178. Gu, Z. (2022). Complex heatmap visualization. Imeta 1, e43. 10.1002/imt2.43.

179. Team, R.C. (2024). R: A Language and Environment for Statistical Computing (R Foundation for Statistical Computing Vienna, Austria).

180. Wickham, H., Averick, M., Bryan, J., Chang, W., McGowan, L.D.A., François, R., Grolemund, G., Hayes, A., Henry, L., Hester, J., et al. (2019). Welcome to the tidyverse. 4, 1686. 10.21105/joss.01686.

181. Fernandez, C.A., Des Rosiers, C., Previs, S.F., David, F., and Brunengraber, H. (1996). Correction of 13C mass isotopomer distributions for natural stable isotope abundance. J Mass Spectrom 31, 255–262. 10.1002/(SICI)1096-9888(199603)31:3<255::AID-JMS290>3.0.CO;2-3.

182. Zougman, A., Selby, P.J., and Banks, R.E. (2014). Suspension trapping (STrap) sample preparation method for bottom-up proteomics analysis. Proteomics 14, 1006–1000. 10.1002/pmic.201300553.

183. Tyanova, S., Temu, T., and Cox, J. (2016). The MaxQuant computational platform for mass spectrometry-based shotgun proteomics. Nat Protoc 11, 2301–2319. 10.1038/nprot.2016.136.

184. Bielow, C., Mastrobuoni, G., and Kempa, S. (2016). Proteomics Quality Control: Quality Control Software for MaxQuant Results. J Proteome Res 15, 777–787. 10.1021/acs.jproteome.5b00780.

185. Bruderer, R., Bernhardt, O.M., Gandhi, T., Miladinovic, S.M., Cheng, L.Y., Messner, S., Ehrenberger, T., Zanotelli, V., Butscheid, Y., Escher, C., et al. (2015). Extending the limits of quantitative proteome profiling with data-independent acquisition and application to acetaminophen-treated three-dimensional liver microtissues. Mol Cell Proteomics 14, 1400–1410. 10.1074/mcp.M114.044305.

