## Supplemental Figures for "Biosynthetic plasticity enables CD8+ T cell functional resilience under nutrient stress"

**Figure S1: Supplemental data for metabolic, regulatory, and functional responses to tumor supernatant and nutrient stress media**

A) Viability and production of TNFa, IFNg, and GrzB in CD8+ T cells treated with tumor-conditioned medium for 6 hours (top) or 24 hours (bottom)

B) Expression of amino acid transporter transcripts after 6 hours of treatment with tumor-conditioned medium

C) Relative amino acid abundance in CD8+ T cells treated with nutrient stress media over 6 hours (top), additionally normalized to control at each time point (bottom).

D) Viability of CD8+ T cells treated with nutrient stress media for 6 hours (top) or 24 hours (bottom)

E) Schematic for kinetics of stress response and stress adaptation fates in CD8+ T cells under nutrient stress

**Figure S2: Supplemental data for transcriptome and translatome analysis**

A) Composition of transcriptome (left) and translatome (right) by transcript type as a % of all transcripts detected

B) Split violin plot of normalized expression values in Control samples by RNA type in input (left) and polysome-bound RNA

C) Heatmap of differentially expressed lon-noncoding RNAs (lncRNAs) in CD8+ T cells under acute nutrient stress

D) Composition of the input RNA and polysome-bound pools by pathway

E) Principal Component Analysis (PCA) of transcript expression level in nutrient stress conditions across whole RNA dataset (left), input RNA (middle) or polysome-bound RNA (right)

F) Rank-ordered plot of gene frequency within the polysome-bound RNA pool in control and No Glutamine conditions. Genes related to T cell function, glycolysis, translation, and stress are highlighted.

G) Upset plots of differentially expressed transcripts in input RNA or polysome-bound RNA pools across nutrient stress conditions. Dots represent conditions where a transcript is differentially expressed in a given direction. Each set is mutually exclusive for a given direction (a transcript can be present in only one set in each direction). Inset shows total number of differentially expressed genes in both up and down directions for each stress condition. DEG = differentially expressed gene / transcript

H) Change in ribosomal and mitochondrial gene expression in nutrient stress conditions vs control for input RNA (left) or polysome-associated RNA (right). All transcripts in the pathway are represented regardless of differential expression status. Violin plot represents distribution and mean of Log2 fold-changes of all transcripts in the pathway, while barcode plot represents Log2 fold-change for each transcript.

I) Change in expression of all transcripts containing a internal ribosomal entry site (IRES) from Thoreen et al. (2012) in input (left) and polysome-associated RNA (right)

**Figure S3: Translation analysis and proteome of CD8+ T cells under nutrient stress**

A) Frequency of transcripts by mode of regulation across nutrient stress conditions according to Anota2seq analysis

B) Scatterplots of change in transcript expression in input (x-axis) and polysome-associated (y-axis) RNA pools across nutrient stress conditions vs. control according to Anota2seq

C) Scatterplot of z-scores of Log2 fold change (stress vs control) for polysome-bound RNA transcripts and corresponding proteins

**Figure S4: Kinetics and dose-dependence of response to nutrient stresses**

A) Visualization of control protein targets (total S6, beta-actin) over time in CD8+ T cells cultured in nutrient stress media.

B) Kinetics of ISR-related transcript induction over time in CD8+ T cells cultured in nutrient stress media.

C) Dose-response of ISR-related transcript induction at 3 hours in nutrient stress media containing varying levels of glucose, glutamine, and methionine. Plotted relative to “1x” concentration in each respective medium (10mM glucose, 2mM glutamine, 100uM methionine, similar to control RPMI medium)

D) ISR-related transcript induction in CD8+ T cells cultured in tumor-conditioned medium for 3 hours

E) Visualization of mTOR and ISR signaling in CD8+ T cells cultured in tumor conditioned medium for 6 hours.

**Figure S5: Contribution of ISR vs mTOR/translation to stress adaptation and programming**
A) Visualization of total S6 and beta-actin in CD8+ T cells cultured in halofuginone, Torin, or 4EGi-1.
B) Kinetics of ISR-related transcript induction in CD8+ T cells cultured in halofuginone, Torin, or 4EGi-1.
C) Viability and production of TNFa, IFNg, and GrzB in CD8+ T cells treated with Torin or 4EGi-1 for 6 hours (top) or 24 hours (bottom).
D) Viability and production of TNFa, IFNg, and GrzB in CD8+ T cells treated with halofuginone for 6 hours (top) or 24 hours (bottom).
E) Representative plots of intracellular cytokine production and surface CD98 expression in CD8+ T cells cultured in halofuginone for 6 hours (top) or 24 hours (bottom)
F) Kinetics of amino acid transporter transcript expression in CD8+ T cells cultured in halofuginone, Torin, or 4EGi-1.
G) Expression of selected amino acid transporters and synthesis enzymes in input (left) and polysome-associated RNA (right).

**Figure S6: GCN2 activity promotes stress resilience in CD8+ T cells**
A) Schematic of the integrated stress response downstream of amino acid stress or halofuginone, highlighting the contribution of GCN2-mediated sensing, translational responses, and transcriptional response downstream of stress sensitive transcription factors.
B) Validation of inhibition of halofuginone-induced ISR signaling by GCN2iB. All samples run on single gel - unused lanes were removed where indicated.
C) Viability and CD98 expression of CD8+ T cells treated with nutrient stress conditions in the presence of DMSO (closed) or GCN2iB (open) for 24 hours.
D) Viability and intracellular cytokine production of CD8+ T cells treated with halofuginone in the presence of DMSO (closed) or GCN2iB (open) for 24 hours.

**Figure S7: ATF4 and CEBPG KO promotes stress amplification and dysfunctional programming**
A) gProfiler2 enrichment plot of pathways significantly enriched in sets of differentially expressed transcripts from Cluster 1-3 (left), Cluster 4 (middle), and Cluster 5 (right). Selected terms are highlighted in tables below.
B) Heatmaps of changes in transcript expression for all annotated genes from indicated pathways.
C) Visualization of ATF4 and p-eIF2a levels in *Rosa26* (control), ATF4 KO, or CEBPG KO CD8+ T cells in the presence of DMSO or halofuginone for 24 hours.
D) Heatmap of differentially expressed transcription factors.

**Figure S8: Additional characterization of stress-sensitive metabolic phenotypes using stable isotope labeling**
A) Quantification of metabolic parameters from Seahorse glycolytic and mitochondrial stress tests in Fig 4I-J.
B) Labeling of additional glycolytic metabolites from U-13C glucose over final 18 hours in *Rosa26* (control), ATF4 KO, or CEBPG KO CD8+ T cells in the presence of DMSO or halofuginone for 24 hours.
C) Labeling of glycine and alanine (m/z 232) from U-13C glucose over final 18 hours in *Rosa26* (control), ATF4 KO, or CEBPG KO CD8+ T cells in the presence of DMSO or halofuginone for 24 hours. Note 1C lost in ionization.
D) Labeling of TCA cycle-related metabolites from U-13C glucose over final 18 hours in *Rosa26* (control), ATF4 KO, or CEBPG KO CD8+ T cells in the presence of DMSO or halofuginone for 24 hours.
E) Labeling of TCA cycle metabolites from U-13C glutamine over final 18 hours in *Rosa26* (control), ATF4 KO, or CEBPG KO CD8+ T cells in the presence of DMSO or halofuginone for 24 hours.
F) Total pool size of leucine and isoleucine in *Rosa26* (control), ATF4 KO, or CEBPG KO CD8+ T cells cultured in the presence of DMSO or halofuginone for 24 hours.

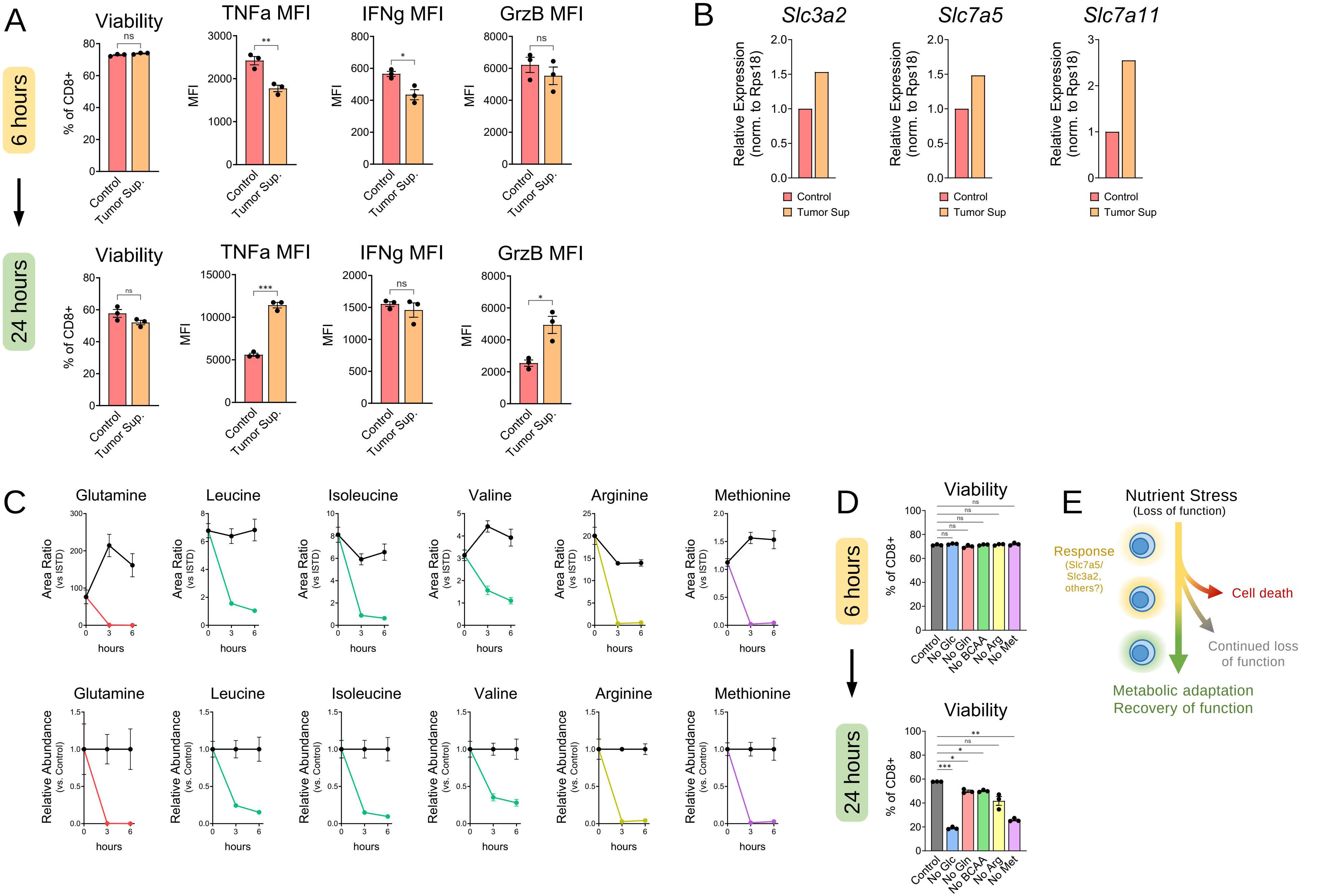
A
6 hours
24 hours
Viability
TNFa MFI
IFNg MFI
GrzB MFI
% of CD8+
MFI
Control
Tumor Sup.
ns
**
*
***
B
Slc3a2
Slc7a5
Slc7a11
Relative Expression (norm. to Rps18)
Control
Tumor Sup
C
Glutamine
Leucine
Isoleucine
Valine
Arginine
Methionine
Area Ratio (vs ISTD)
Relative Abundance (vs. Control)
hours
D
Viability
No Glc
No Gln
No BCAA
No Arg
No Met
E
Nutrient Stress
(Loss of function)
Response
(Slc7a5/ Slc3a2, others?)
Cell death
Continued loss of function
Metabolic adaptation
Recovery of function

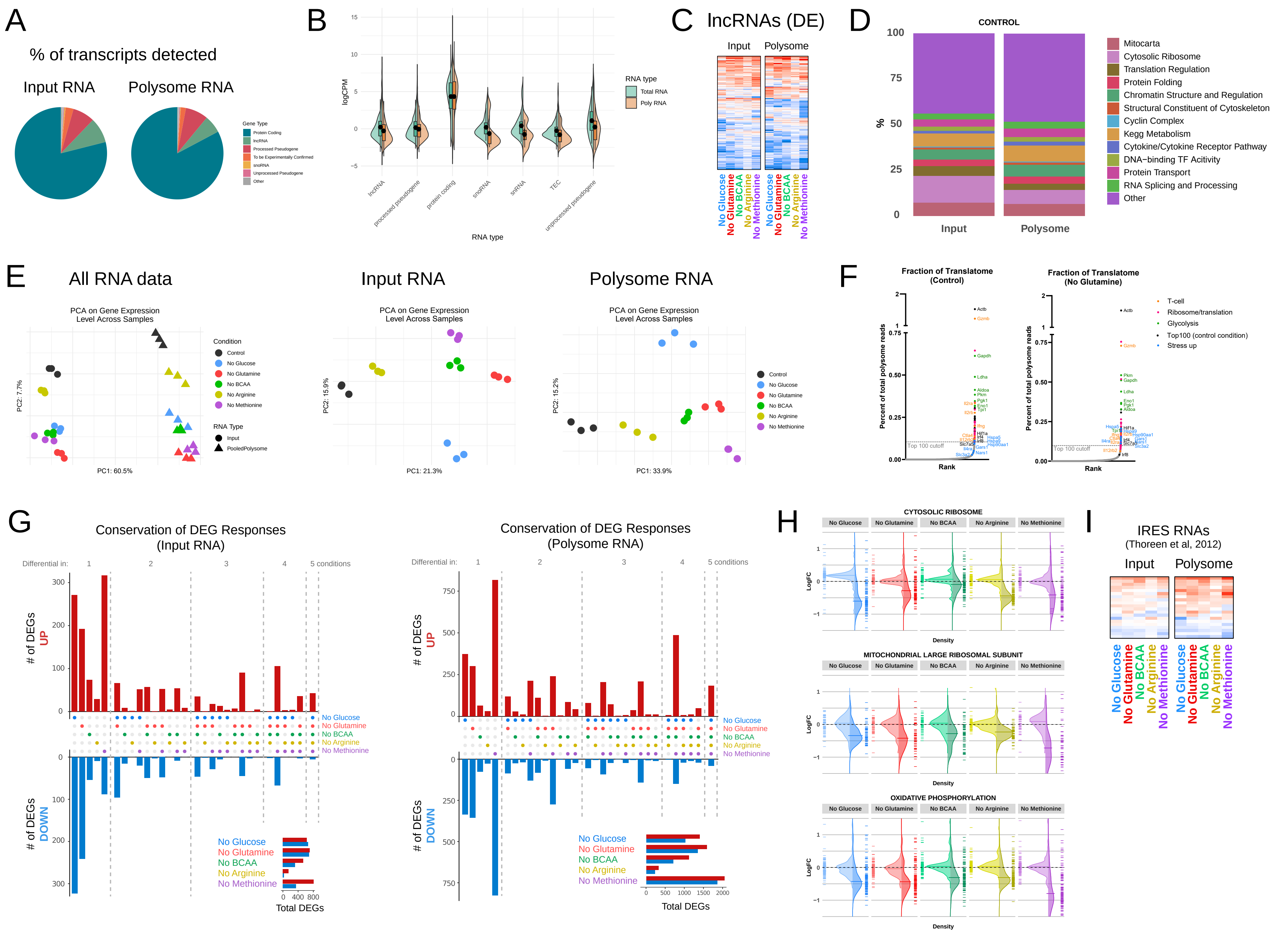
A
% of transcripts detected
Input RNA
Polysome RNA
Gene Type
Protein Coding
lncRNA
Processed Pseudogene
To be Experimentally Confirmed
snoRNA
Unprocessed Pseudogene
Other
B
logCPM
RNA type
Total RNA
Poly RNA
lncRNA
processed pseudogene
protein coding
snoRNA
snRNA
TEC
unprocessed pseudogene
C
lncRNAs (DE)
Input
Polysome
No Glucose
No Glutamine
No BCAA
No Arginine
No Methionine
D
CONTROL
%
Input
Polysome
Mitocarta
Cytosolic Ribosome
Translation Regulation
Protein Folding
Chromatin Structure and Regulation
Structural Constituent of Cytoskeleton
Cyclin Complex
Kegg Metabolism
Cytokine/Cytokine Receptor Pathway
DNA−binding TF Acitivity
Protein Transport
RNA Splicing and Processing
Other
E
All RNA data
Input RNA
Polysome RNA
PCA on Gene Expression Level Across Samples
PC1: 60.5%
PC2: 7.7%
PC1: 21.3%
PC2: 15.9%
PC1: 33.9%
PC2: 15.2%
Condition
Control
No Glucose
No Glutamine
No BCAA
No Arginine
No Methionine
RNA Type
Input
PooledPolysome
F
Fraction of Translatome (Control)
Fraction of Translatome (No Glutamine)
Percent of total polysome reads
Rank
Top 100 cutoff
T-cell
Ribosome/translation
Glycolysis
Top100 (control condition)
Stress up
G
Conservation of DEG Responses (Input RNA)
Conservation of DEG Responses (Polysome RNA)
Differential in:
5 conditions
### of DEGs UP
### of DEGs DOWN
Total DEGs
H
CYTOSOLIC RIBOSOME
MITOCHONDRIAL LARGE RIBOSOMAL SUBUNIT
OXIDATIVE PHOSPHORYLATION
LogFC
Density
I
IRES RNAs
(Thoreen et al, 2012)
Input
Polysome

Supplemental Figure 3

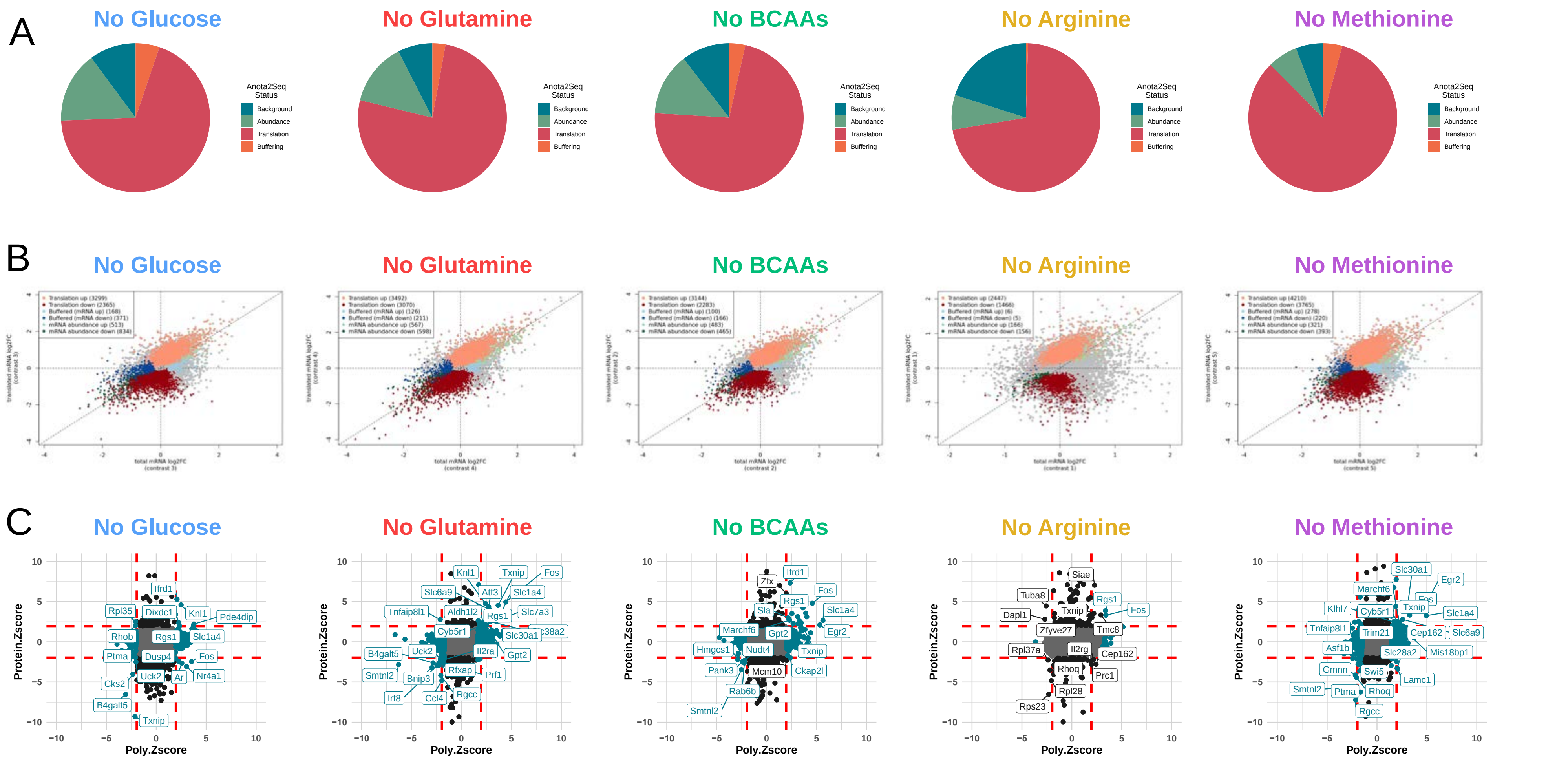

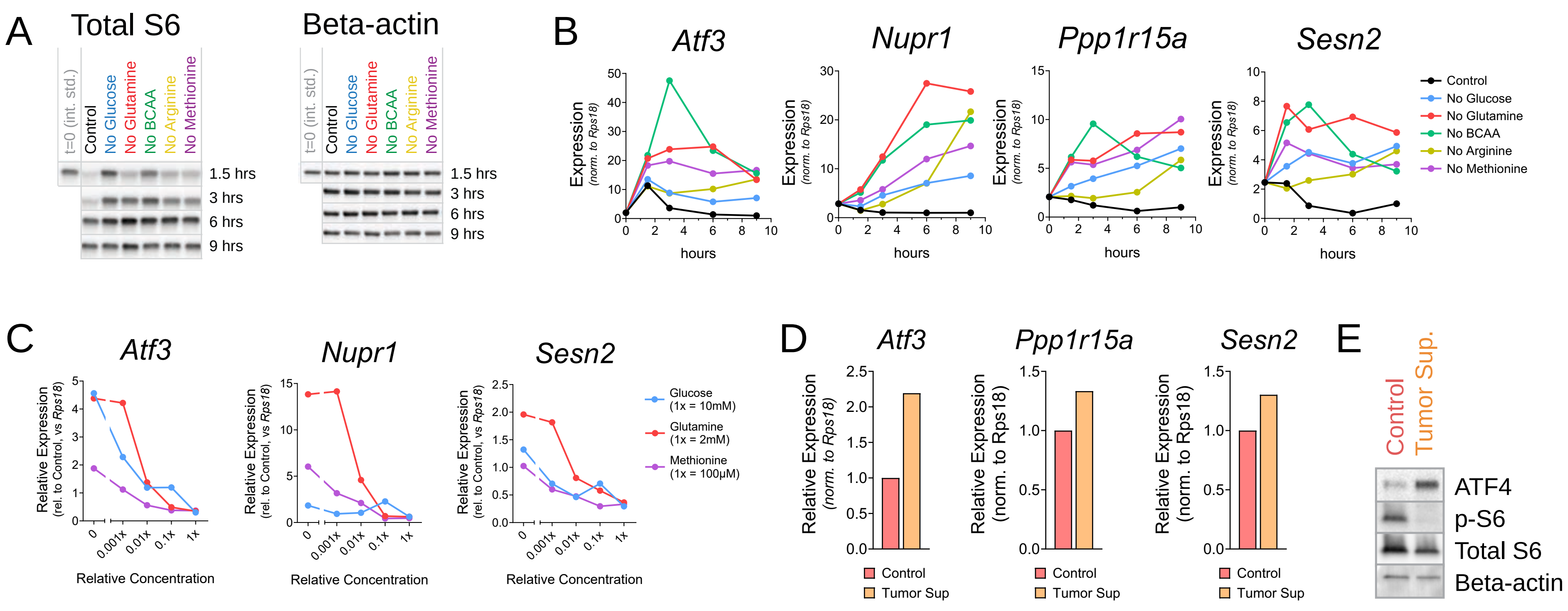
A
Total S6
Beta-actin
t=0 (int. std.)
Control
No Glucose
No Glutamine
No BCAA
No Arginine
No Methionine
1.5 hrs
3 hrs
6 hrs
9 hrs
B
Atf3
Nupr1
Ppp1r15a
Sesn2
Expression (norm. to Rps18)
hours
Control
No Glucose
No Glutamine
No BCAA
No Arginine
No Methionine
C
Atf3
Nupr1
Sesn2
Relative Expression (rel. to Control, vs Rps18)
Relative Concentration
0
0.001x
0.01x
0.1x
1x
Glucose (1x = 10mM)
Glutamine (1x = 2mM)
Methionine (1x = 100µM)
D
Atf3
Ppp1r15a
Sesn2
Relative Expression (norm. to Rps18)
Control
Tumor Sup
E
Control
Tumor Sup.
ATF4
p-S6
Total S6
Beta-actin

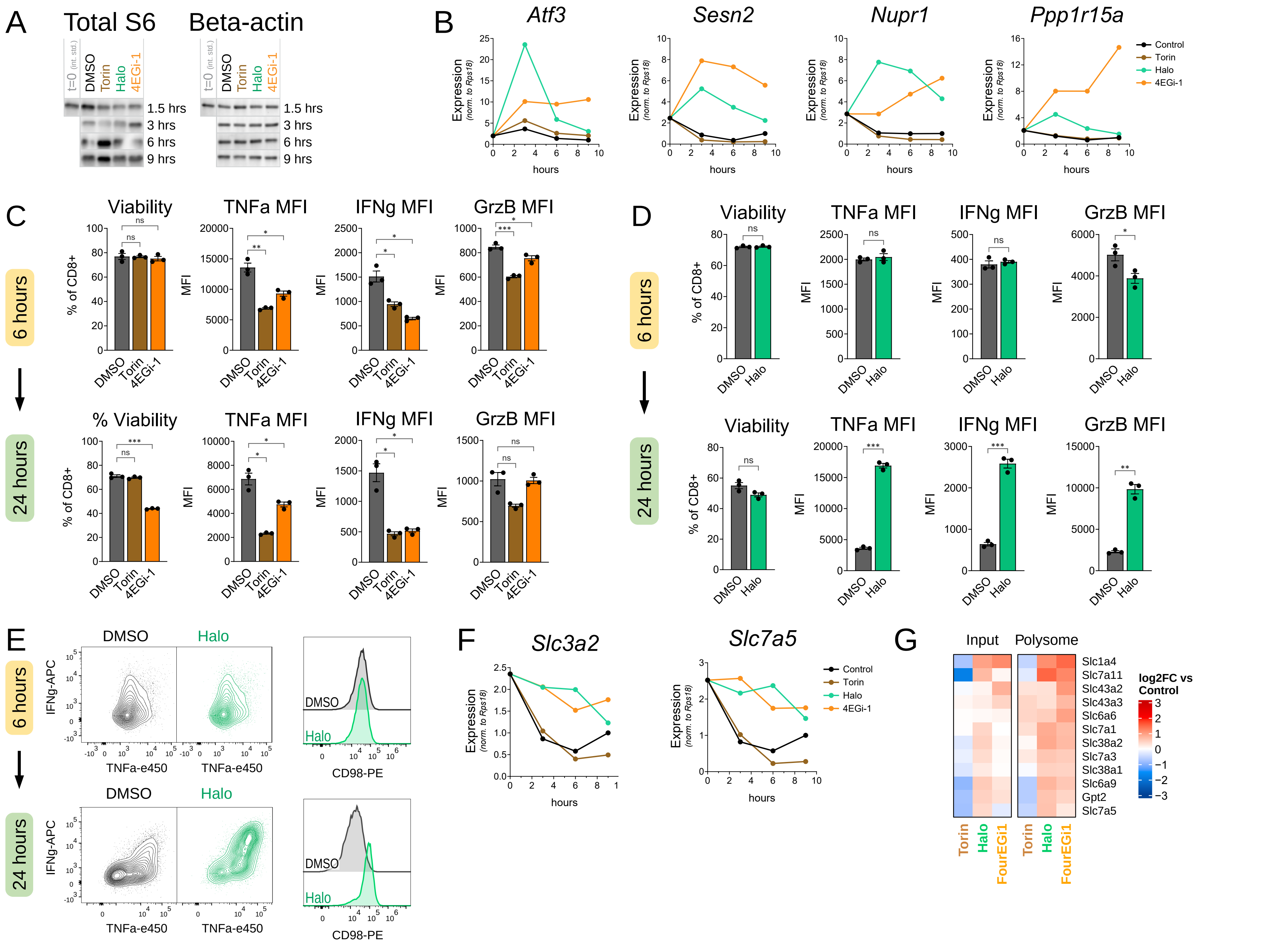
A
Total S6
Beta-actin
t=0 (int. std.)
DMSO
Torin
Halo
4EGi-1
1.5 hrs
3 hrs
6 hrs
9 hrs
B
Atf3
Sesn2
Nupr1
Ppp1r15a
Expression (norm. to Rps18)
hours
Control
Torin
Halo
4EGi-1
C
D
6 hours
24 hours
Viability
% Viability
TNFa MFI
IFNg MFI
GrzB MFI
% of CD8+
MFI
DMSO
Torin
4EGi-1
Halo
ns
E
DMSO
Halo
IFNg-APC
TNFa-e450
CD98-PE
F
Slc3a2
Slc7a5
G
Input
Polysome
Slc1a4
Slc7a11
Slc43a2
Slc43a3
Slc6a6
Slc7a1
Slc38a2
Slc7a3
Slc38a1
Slc6a9
Gpt2
Slc7a5
log2FC vs Control
FourEGi1

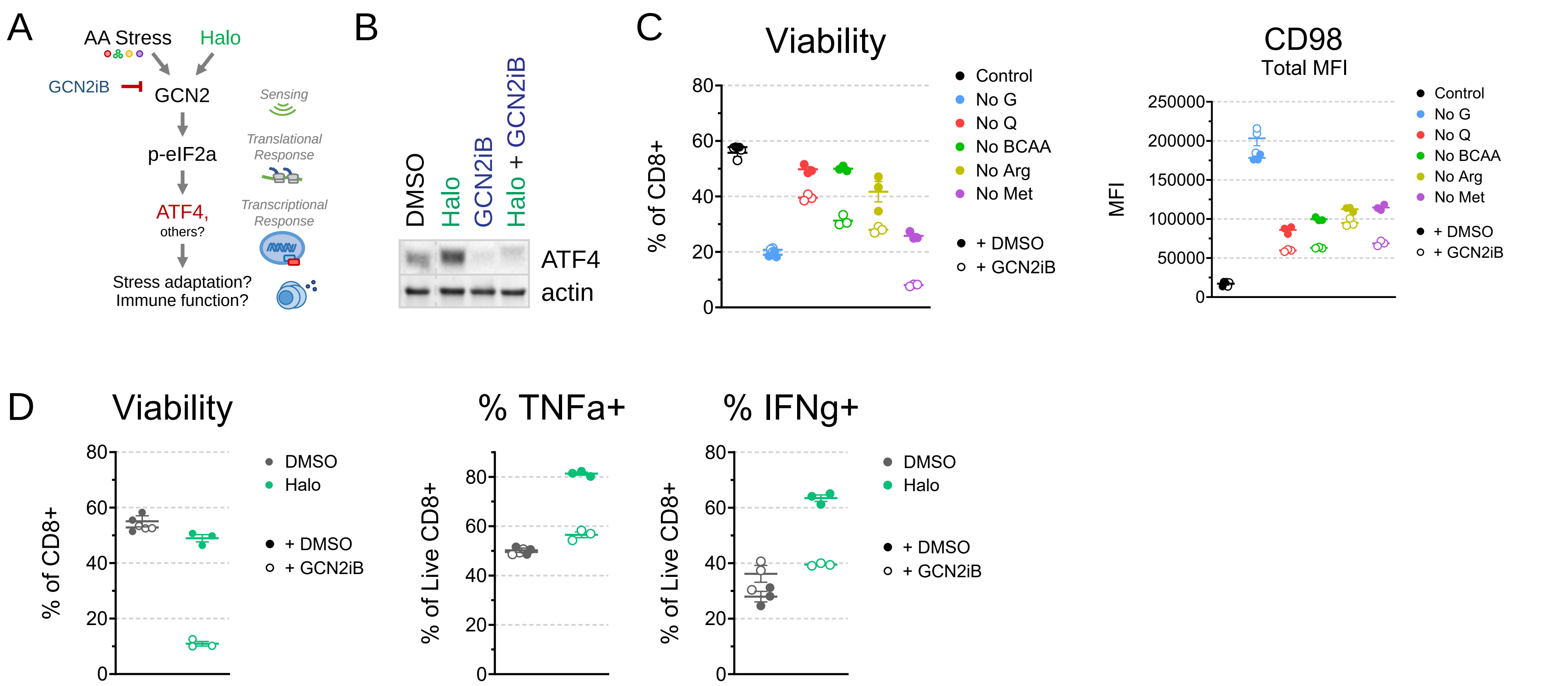
A
AA Stress
Halo
GCN2iB
GCN2
Sensing
p-eIF2a
Translational Response
ATF4,
others?
Transcriptional Response
Stress adaptation?
Immune function?
B
DMSO
Halo
GCN2iB
Halo + GCN2iB
ATF4
actin
C
Viability
% of CD8+
Control
No G
No Q
No BCAA
No Arg
No Met
+ DMSO
+ GCN2iB
CD98
Total MFI
MFI
D
Viability
% of CD8+
DMSO
Halo
+ DMSO
+ GCN2iB
% TNFa+
% of Live CD8+
% IFNg+
% of Live CD8+

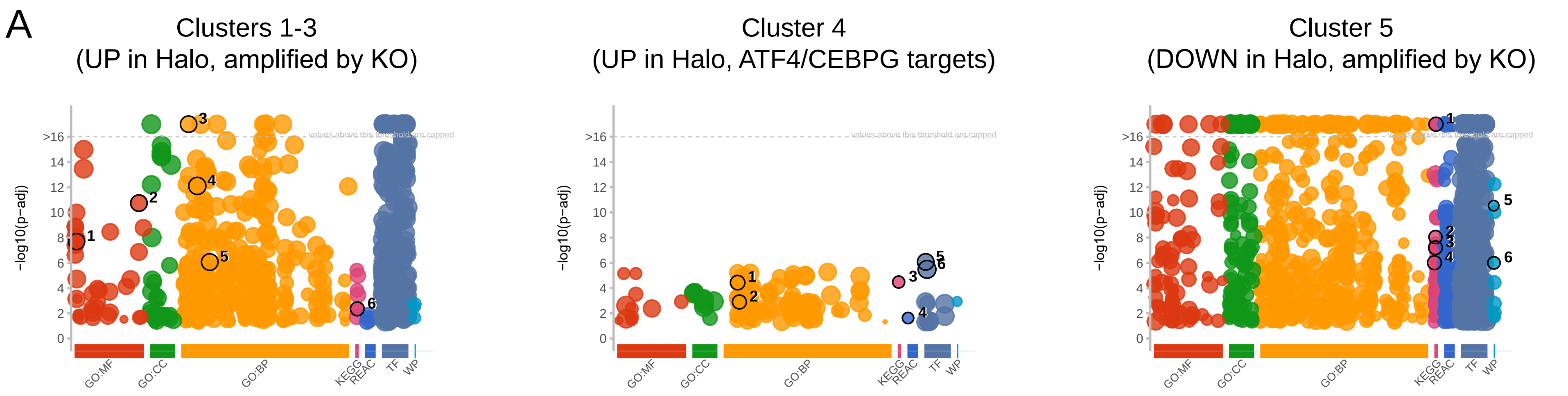


| id | source | term_id | term_name | term_size | | p_value |
|---|---|---|---|---|---|---|
| 1 | GO:MF | GO:0003700 | DNA−binding transcription factor activity | 1239 | | 2.0e−08 |
| 2 | GO:MF | GO:0140110 | transcription regulator activity | 1655 | | 1.8e−11 |
| 3 | GO:BP | GO:0002682 | regulation of immune system process | 1502 | | 2.0e−17 |
| 4 | GO:BP | GO:0006950 | response to stress | 3385 | | 7.8e−13 |
| 5 | GO:BP | GO:0012501 | programmed cell death | 1826 | | 8.6e−07 |
| 6 | KEGG | KEGG:04630 | JAK−STAT signaling pathway | 171 | | 4.4e−03 |

g:Profiler (biit.cs.ut.ee/gprofiler)

| id | source | term_id | term_name | term_size | | p_value |
|---|---|---|---|---|---|---|
| 1 | GO:BP | GO:0006520 | amino acid metabolic process | 232 | | 3.7e−05 |
| 2 | GO:BP | GO:0006865 | amino acid transport | 154 | | 1.3e−03 |
| 3 | KEGG | KEGG:00970 | Aminoacyl−tRNA biosynthesis | 44 | | 3.3e−05 |
| 4 | REAC | REAC:R−MMU−352230 | Amino acid transport across the plasma membrane | 29 | | 2.4e−02 |
| 5 | TF | TF:M10181 | Factor: ATF−4; motif: RNMTGATGCAAY | 1304 | | 8.4e−07 |
| 6 | TF | TF:M10196 | Factor: C/EBPgamma; motif: NNMTGATGCAAY | 4665 | | 3.3e−06 |

g:Profiler (biit.cs.ut.ee/gprofiler)

| id | source | term_id | term_name | term_size | | p_value |
|---|---|---|---|---|---|---|
| 1 | KEGG | KEGG:04110 | Cell cycle | 155 | | 1.4e−27 |
| 2 | KEGG | KEGG:01230 | Biosynthesis of amino acids | 78 | | 9.0e−09 |
| 3 | KEGG | KEGG:01200 | Carbon metabolism | 120 | | 6.3e−08 |
| 4 | KEGG | KEGG:00190 | Oxidative phosphorylation | 134 | | 9.7e−07 |
| 5 | WP | WP:WP103 | Cholesterol biosynthesis | 15 | | 2.9e−11 |
| 6 | WP | WP:WP157 | Glycolysis and gluconeogenesis | 49 | | 1.0e−06 |

g:Profiler (biit.cs.ut.ee/gprofiler)

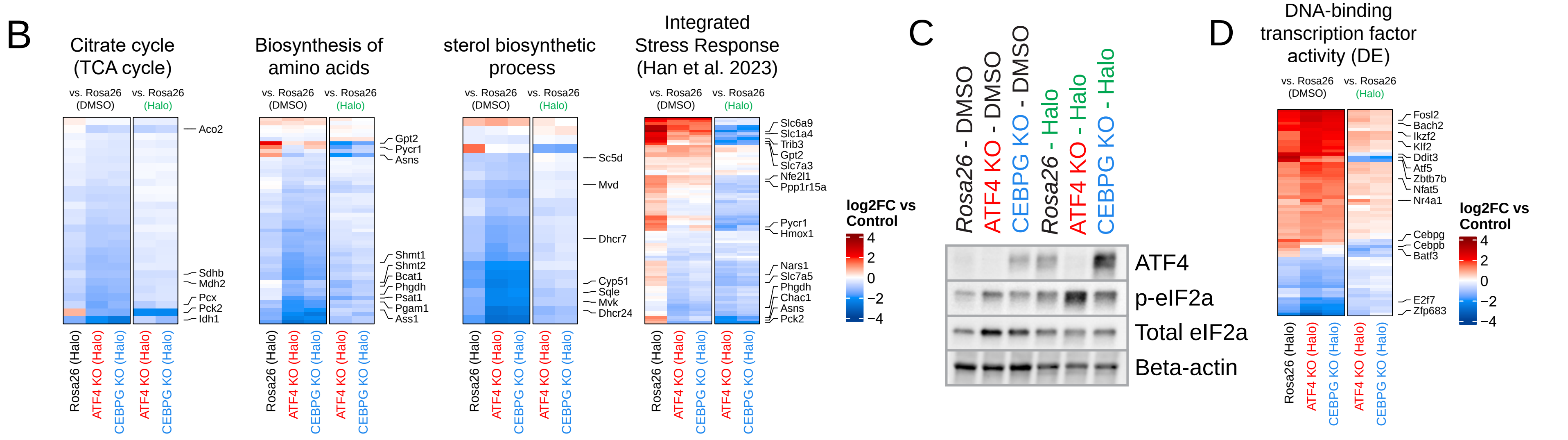

Supplemental Figure 8

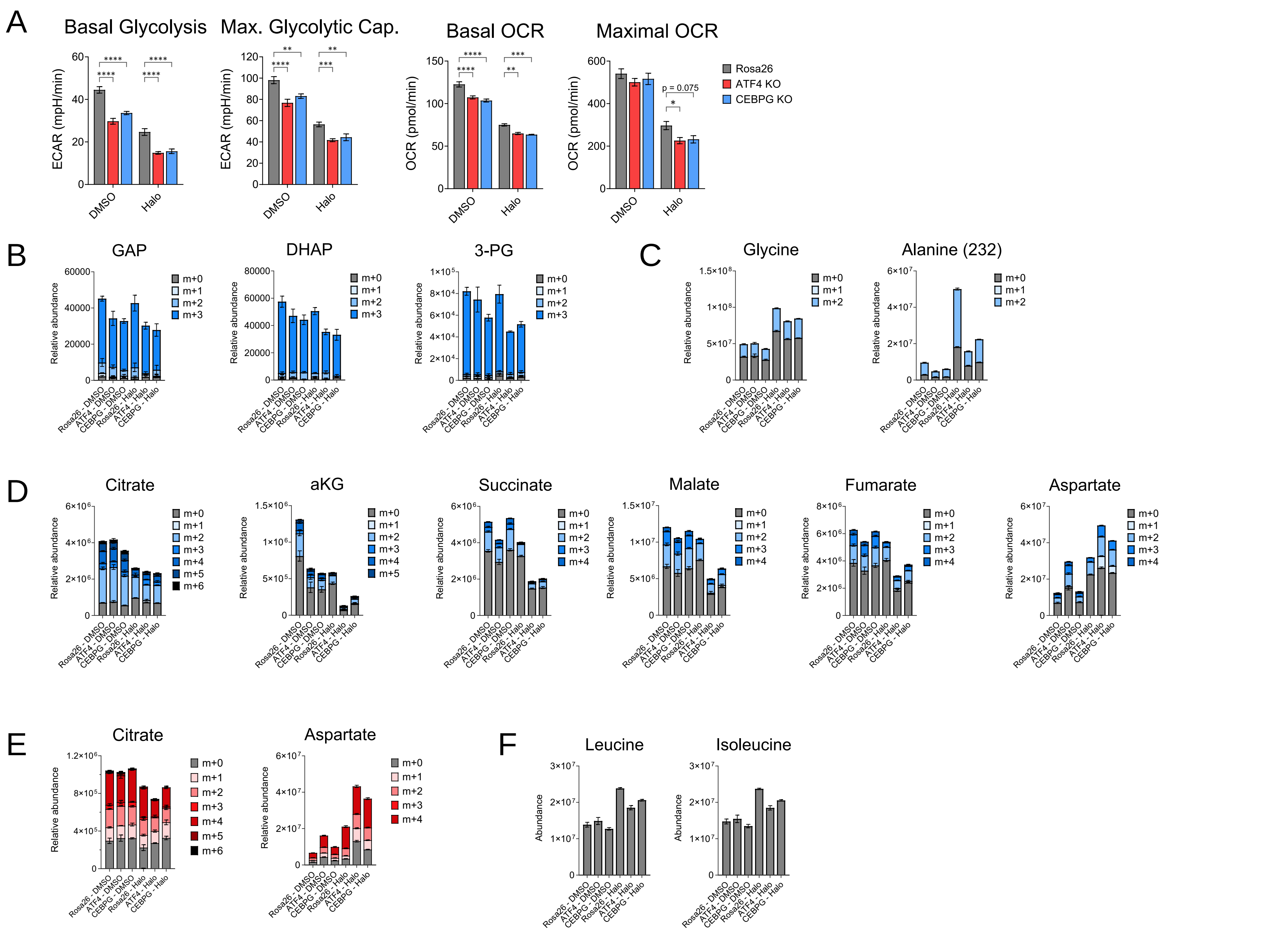

**Table S1: Retention times and m/z for GC-MS analysis of stable isotope labeling, related to STAR Methods**

| Metabolite | Retention time | m/z |
|---|---|---|
| Dihydroxyacetone phosphate | 24.73 | 484 |
| Glyceraldehyde-3-phosphate | 24.96 | 484 |
| 3-phosphoglycerate | 27.22 | 585 |
| Phosphoenolpyruvate | 23.06 | 453 |
| Pyruvate | 7.24 | 174 |
| Lactate | 11.41 | 233/**261** |
| Citrate | 27.12 | 459/**591** |
| a-ketoglutarate | 19.91 | 346 |
| Succinate | 15.99 | 289 |
| Fumarate | 16.44 | 287 |
| Malate | 21.4 | 419 |
| Glycerol-3-phosphate | 24.46/26.83 | 571 |
| Alanine | 12.25 | **232**/**260** |
| Aspartate | 22.01 | 302/390/**418** |
| Glutamate | 26.6 | 330/**432** |
| Glutamine | 25.37 | 431 |
| Glycine | 12.62 | 218/**246** |
| Leucine | 15.00 | 200 |
| Isoleucine | 15.54 | 200 |
| Serine | 19.64 | 288/302/362/**390** |

When multiple m/z were generated, all were used to identify the peaks, but the m/z in bold was used in the quantification for data visualization.
